# A nuclear hormone receptor and lipid metabolism axis are required for the maintenance and regeneration of reproductive organs

**DOI:** 10.1101/279364

**Authors:** Shasha Zhang, Longhua Guo, Carlos Guerrero-Hernández, Eric J Ross, Kirsten Gotting, Sean A. McKinney, Wei Wang, Youbin Xiang, R. Scott Hawley, Alejandro Sánchez Alvarado

## Abstract

Understanding how stem cells and their progeny maintain and regenerate reproductive organs is of fundamental importance. The freshwater planarian *Schmidtea mediterranea* provides an attractive system to study these processes because its hermaphroditic reproductive system (RS) arises post-embryonically and when lost can be fully and functionally regenerated from the proliferation and regulation of experimentally accessible stem and progenitor cells. By controlling the function of a nuclear hormone receptor gene (*nhr-1*), we established conditions in which to study the formation, maintenance and regeneration of both germline and somatic tissues of the planarian RS. We found that *nhr-1*(*RNAi*) not only resulted in the gradual degeneration and complete loss of the adult hermaphroditic RS, but also in the significant downregulation of a large cohort of genes associated with lipid metabolism. One of these, *Smed-acs-1*, a homologue of Acyl-CoA synthetase, was indispensable for the development, maintenance and regeneration of the RS, but not for the homeostasis or regeneration of other somatic tissues. Remarkably, supplementing *nhr-1*(*RNAi*) animals with either bacterial Acyl-CoA synthetase or the lipid metabolite Acetyl-CoA rescued the phenotype restoring the maintenance and function of the hermaphroditic RS. Our findings uncovered a likely evolutionarily conserved role for nuclear hormone receptors and lipid metabolism in the regulation of stem and progenitor cells required for the long-term maintenance and regeneration of animal reproductive organs, tissues and cells.

## INTRODUCTION

The adult organs of most organisms are actively maintained by a complex interplay of cellular and metabolic homeostatic processes that include hormonal regulation, the maintenance of stem cell pools, removal of degenerated cells and the generation and functional integration of new cells through proliferation and differentiation (Guo and Cantley, 2010; Knapp and Tanaka, 2012; Rock and Hogan, 2011; Tu et al., 2016). This plasticity is observed in organs such as lung, muscle, skin, heart, and liver, but it is most dramatically manifested in the reproductive system (RS) of many animals. In humans, many organs of the RS are highly plastic in adulthood and require cyclical regeneration and extensive remodeling (Nair and Taylor, 2010). For instance, the human endometrium undergoes growth, differentiation and shedding during every menstrual cycle, a homeostatic process that requires the coordination of proliferation and differentiation of epithelial progenitor and mesenchymal/stromal stem cells with estrogen and progesterone fluctuations during the estrus cycle (Gargett et al., 2012).

Although it is well-established that nuclear hormone receptors (NHRs) for androgen (Yeh et al., 2002), progesterone (Chappell et al., 1997) and estrogen (Walker and Korach, 2004) are essential for RS development and function, precisely how the endocrine system affects the stem cell populations responsible for the maintenance and cyclical regeneration of adult RS organs remains incompletely understood. For example, the precise location and types of both myometrial and endometrial stem cells chiefly responsible for the 500- to 1,000- fold increase in volume and 24-fold increase in weight of the human uterus have yet to be fully determined (Kørbling and Estrov, 2003; Ono et al., 2008; Ramsey, 1994; Shynlova et al., 2006). However, the deep evolutionary conservation of endocrine regulation of the RS has allowed research organisms such as fruitflies and nematodes to shed light on some of the roles lipophilic hormones play in the regulation of germ cells, and the embryonic development of reproductive organs (Allen and Spradling, 2008; Asahina et al., 2000; Gissendanner et al., 2004; Gissendanner et al., 2008; Sun and Spradling, 2013).

Lipid metabolism also plays an evolutionarily conserved role during RS embryonic development and fertility (Barton et al., 2016; Sano et al., 2005). Cholesterol and β-oxidation of fatty acids are essential for meiosis, embryo development, and uterus function in both mice and humans (Downs et al., 2009; Dunning et al., 2010; Mouzat et al., 2013; Seli et al., 2014). In *D. melanogaster*, female feeding behavior, lipid accumulation in the oocyte, and diet nutrient coordination, lipid metabolism and oocyte development are controlled by the Ecdysone receptor (EcR) (Sieber and Spradling, 2015). And in *Caenorhabditis elegans*, fatty acids and their derivatives not only influence reproductive growth and fertilization (Tang and Han, 2017; Wang et al., 2015; Zhu and Han, 2014), but also regulate germ cell fate via Acyl CoA synthetase and its downstream product Myristoyl-CoA (Tang and Han, 2017).

The fundamental importance of lipophilic hormones and their receptors on sexual reproduction is underscored by the effect metabolism has on endocrine functions. When systemic metabolism fails due to either disease or aging (Conboy and Rando, 2005; Lopez-Otin et al., 2016), both germ and somatic cells in the RS gradually degenerate in both men and women (Makabe et al., 1998; Motta et al., 2002; Paniagua et al., 1991). Changes in fat and energy storage in the body have been shown to affect the reproductive activity of many vertebrates (Ballinger, 1977; Eliassen and Vahl, 1982; Reznick and Braun, 1987). Similarly, the adult females of *D. melanogaster* can remodel their midgut to adjust the energy balance between the body and the RS after mating (Reiff et al., 2015) and in *C. elegans* dietary restriction has been shown to delay aging related degeneration of the egg-laying apparatus (Pickett and Kornfeld, 2013). Conversely, age-related and disease-induced changes in RS metabolism also feed back to the metabolic homeostasis of the body. For example, menopause alters lipid metabolism in adipose tissue causing atopic lipid accumulation in liver, macrophages, and the cardiovascular system (Della Torre et al., 2014); and diseases of the RS, such as polycystic ovary syndrome, are often associated with metabolic ailments such as nonalcoholic fatty liver disease (Della Torre et al., 2014).

While the evolutionarily conserved roles for NHR and lipid metabolism in RS embryonic development and germ stem cell functions are well established, their precise role in regulating somatic stem and progenitor cell functions during adult RS maintenance and regeneration remains an open question. The recent discovery that the nuclear hormone receptor *nhr-1* is required for the post-embryonic development of the hermaphroditic RS in the freshwater planarian *Schmidtea mediterranea* (Tharp et al., 2014), has made this organism an attractive system to study the roles that NHRs and lipid metabolism may play in the plasticity of the RS. Unlike other invertebrate research organisms (*i.e*., fruitflies and nematodes), planarians are devoid of both gonads and attendant somatic organs and tissues of the RS upon birth. As animals grow, the entire RS, including the germline, arises from the proliferation and differentiation of adult stem cells known as neoblasts (Newmark et al., 2008). Once established, the RS may be subjected to changes related to either injury and/or nutritional intake. If an animal is amputated, the resulting fragments resorb their RS as they regenerate the missing body parts. A similar resorption of the RS is also observed when the animals are subjected to starvation. In both cases, once food becomes available, the animals regenerate a fully functional RS (Guo et al., 2016; Newmark and Sanchez Alvarado, 2002).

Numerous evolutionarily conserved genes and pathways have also been shown to regulate RS development, homeostasis and regeneration in *S. mediterranea* (Chong et al., 2013; Guo et al., 2016; Iyer et al., 2016a; Newmark and Sanchez Alvarado, 2002; Newmark et al., 2008). For instance, *Dmd1*, one the most conserved genes in male sex determination across bilaterians, is essential for the maintenance of the planarian male RS, including germ stem cell specification, and differentiation (Chong et al., 2013). *nanos*, the nuclear factor Y-B (*NF-YB*), *Smed-boule2* and several azoospermia (DAZ) family members are also required for germ stem cell maintenance (Iyer et al., 2016a; Iyer et al., 2016b; Wang et al., 2010; Wang et al., 2007). And *Smed-boule1*, homologs of vertebrate DAZ-associated proteins, synaptonemal complex protein 1 (SYCP1), *Smed-CPEB, Smed-eIF4E-like* and t-complex proteins have all been shown to be essential for meiosis in testis (Iyer et al., 2016b; Rouhana et al., 2017; Xiang et al., 2014). Moreover, the size of testes and sperm maturation are determined by the neuropeptide, NPY-8, and its upstream prohormone convertase, *PC2* (Collins et al., 2010), and depletion of insulin-like peptide or insulin receptor have been shown to affect spermatogenesis (Miller and Newmark, 2012). Therefore, *S. mediterranea*, an animal in which the molecular and cellular interactions underpinning the maintenance and regeneration of the RS can be mechanistically dissected, provides unique opportunities to advance our understanding of reproductive biology.

Here, we exploit the biology, evolutionary conservation of regulatory networks and the functional genomic tools available in *S. mediterranea* to uncover molecular mechanisms driving the maintenance and regeneration of the RS. We carried out a discrete RNA mediated genetic interference (RNAi) screen of genes found to be preferentially expressed in sexually mature animals, and found new specific functions in the maintenance and regeneration of the planarian hermaphroditic RS for the previously identified NHR *nhr-1* (Tharp et al., 2014). We report that *nhr-1*(*RNAi*) not only resulted in the gradual degeneration and complete loss of the hermaphroditic RS, but also in the significant downregulation of a cohort of novel reproductive accessory gland markers and a large set of genes associated with lipid metabolism. We discovered that the planarian gene *Smed-acs-1*, a homologue of Acyl-CoA synthetase (ACS), is indispensable for the development, maintenance and regeneration of RS organs and tissues, but not for the homeostasis or regeneration of other somatic tissues. Remarkably, supplementing *nhr-1*(*RNAi*) animals with either bacterial ACS or the lipid metabolite Acetyl-CoA rescued the phenotype restoring the maintenance and function of the hermaphroditic RS. Our findings not only demonstrate that the planarian RS depends on an NHR/lipid metabolism axis for its maintenance and regeneration, but also may have uncovered a conserved molecular mechanism regulating reproductive capacity in long-lived organisms.

## RESULTS

### *nhr-1* is required for the homeostatic maintenance of adult sexual reproductive organs

We sought to identify genes important for the maintenance and regeneration of the RS in *S. mediterranea* by comparing the transcriptomes of decapitated sexually mature animals to immature juveniles. 7,154 transcripts (7.89% Figure 1A) were found to be highly expressed in sexually mature animals. We prioritized for study genes with coiled-coil domains or zinc finger domains (n=34), because proteins with these domains are among the most commonly associated with biological functions in meiosis, hormonal response, and RS development in eukaryotic organisms (Lupas and Bassler, 2017; Razin et al., 2012; Truebestein and Leonard, 2016). Whole mount *in situ* hybridizations confirmed that 30 genes were expressed in the gonads (88.2%, Figure S1A and S1B), and 10 genes (29.4%, Figure S1A and S1B) were expressed in the accessory reproductive organs. Only 2 genes (5.9% Fig S1A and S1B) did not have detectable expression in the RS. The high levels of expression of genes coding for proteins with coiled-coil or zinc finger domains in RS suggested that these proteins may be important for RS functions.

**Figure 1:**
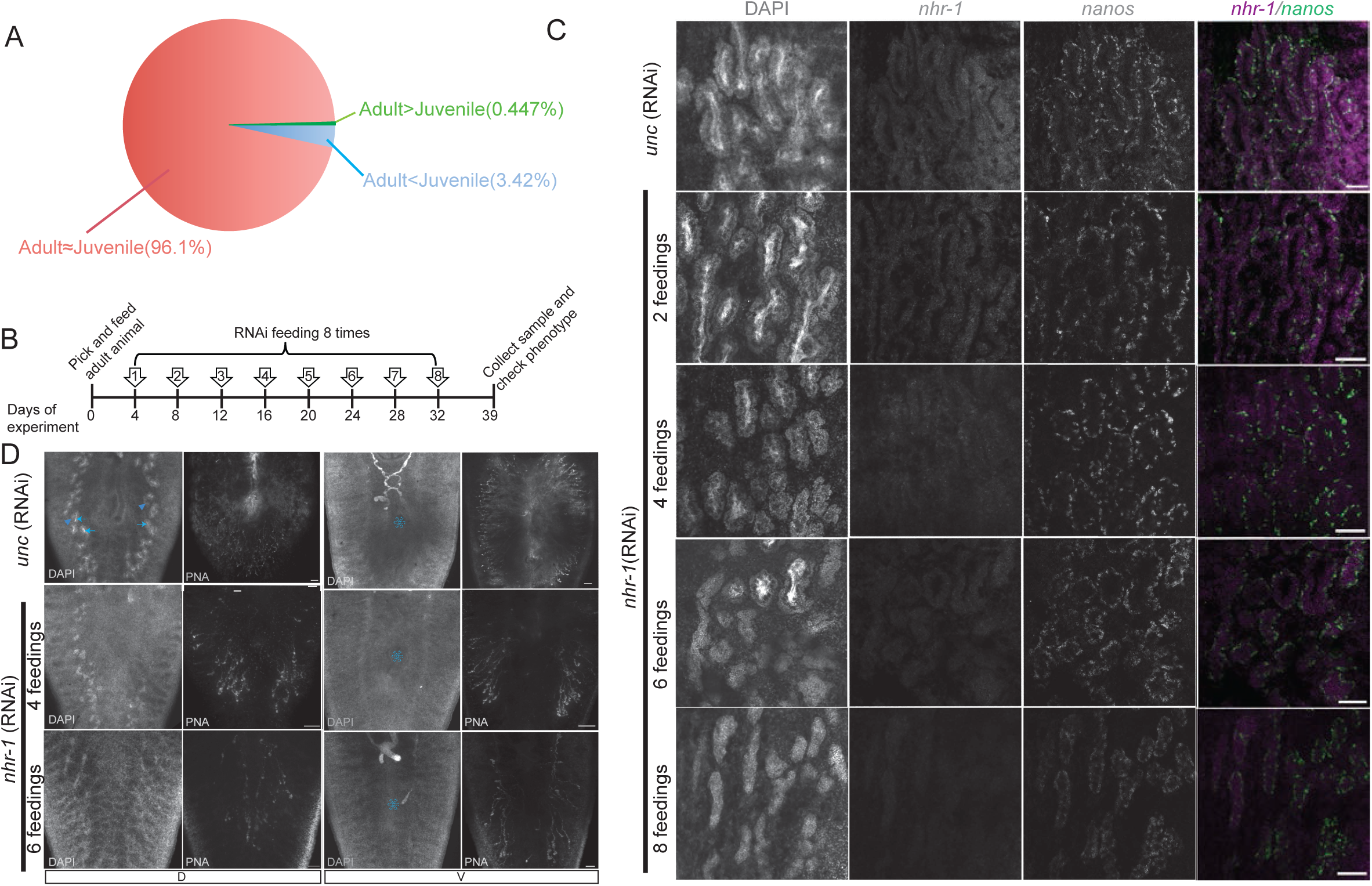
Functional screen of sexual planarian enriched genes revealed *nhr-1* as an essential gene for reproductive system maintenance. (A) Comparison of the juvenile and sexual adult planarian transcriptome (n=3 replicates, each containing 3 planarians. “D0” represents decapitated adult animals. 7.89% transcripts were expressed significantly higher in adult animals (in green, t-test, adj. p<0.05), while 2.58% of transcripts were expressed higher in juvenile animals (in blue, t-test, adj. p<0.05). (B) RNAi feeding schedule diagram. Adult planarians were given RNAi by feeding 8 times as indicated and checked 7 days after the last round of feeding. (C) *In situ* test of *nhr-1* and *nanos* expression in testes. DAPI staining showed the observed testes morphology at the same anatomical location. Scale bar: 100µm (D) Nuclear and gland staining of the *nhr-1*(*RNAi*) animals. DAPI staining was stronger in spermatids (blue arrowheads), spermatozoa and other testes cells (blue triangles) compared to somatic tissues. Penis papilla was also visible by DAPI staining (blue asterisks). Glands were stained by PNA. Scale bar: 20µm

Next, we tested the functions of the identified genes in sexually mature animals by RNAi, followed by DAPI and peanut agglutinin (PNA) staining (Figure 1B). Among the genes found to be essential for RS maintenance was the recently described nuclear hormone receptor *nhr-1*, which was shown to be required for the post-embryonic development of the planarian RS (Tharp et al., 2014). *nhr-1* possesses two zinc finger domains of the C4 class near its N terminus (PFAM Bit scores 38.0 and 92.6, respectively). We tested the function of this receptor in the mature RS by subjecting animals to multiple rounds of RNAi treatment (Figure 1B). Initially, 100% of testes had mature spermatids, but after 4 rounds of treatment 29.4% of testes (5 out of 17) had lost mature spermatids. By 8 rounds of treatment, spermatids could not be detected in 100% of testes assayed (Figure 1C). The accessory reproductive glands can be detected by PNA staining (Chong et al., 2011) in both dorsal and ventral sides of sexually mature animals. PNA staining showed that the reproductive glands started to degenerate after 4 rounds of RNAi feedings and completely disappeared after 8 rounds of treatment (Figure 1D). Interestingly, while *nhr-1*(*RNAi*) did not lead to significant changes in the expression of *nanos* in germ stem cells, even after prolonged RNAi feeding (Figure 1C), *nhr-1*(*RNAi*) worms were sterile, and did not lay any egg capsules (Figure S2). Knockdown of a meiosis specific gene (*sycp1*), or an ovary related gene (*gld*) did not affect fertility (Figure S2). These results indicate that *nhr-1* is essential for the homeostatic maintenance of differentiated germ cells and somatic tissues of the RS.

### *nhr-1* is required for the progression of normal meiosis in testes

We sought to better characterize the observed defect in male meiosis caused by of *nhr-1*(*RNAi*). We quantified the percentage of male germ cells at different stages of meiosis in the testes of both control and *nhr-1*(*RNAi*) animals using DAPI and two-photon confocal microscopy. Multiple testes from the same locations of different animals were imaged from dorsal surface of testes to the middle of the testes at an approximate depth of 50µm, which was sufficient to consistently distinguish male germ cells at different meiotic stages based on their nuclear morphology (Movie-S1). Knocking down *nhr-1* led to decreased germ cells at pachytene stages and to a reduced number of round spermatids (Movie-S2, Figure S3A). The distribution of spermatocytes at the pachytene stage was also decreased (Figure 2B). Since bouquet disruption changes the distribution of prophase cells (Chretien, 2011), we also checked bouquet formation in the spermatocytes (Figures 2A, 2C and S3B). During male meiosis, the telomeres of all 8 chromosomes cluster at the nuclear envelope in the early leptotene stage to form a bouquet, which is considered essential for the progression of meiosis (Figures 2C and S3B)(Chretien, 2011; Xiang et al., 2014). The bouquets persist until the pachytene stage (Figure 2C). In normal male meiosis, 50% of the spermatocytes at leptotene stage had bouquets (Figure 2C). Almost 100% of the spermatocytes at zygotene or pachytene stages formed bouquets (79.2% in zygotene stage and 94% in pachytene stage; Figure 2C). In *nhr-1 RNAi* animals, even though chromosome condensation at each prophase stages was similar to regular meiosis, the telomeres were scattered around or clustered in several locations of the nuclear envelope (Figure 2C). After 6 rounds of *nhr-1*(*RNAi*) treatment, spermatocytes at leptotene and zygotene stages had significantly fewer bouquets (33.3% in leptotene stage, 52.8% in zygotene stage and 84.6% in pachytene stage. For leptotene stage, p=0.0034; for zygotene stage p<0.0001; for pachytene stage, p=0.172. Figure 2A). These results indicate that in the absence of *nhr-1* function, bouquet formation is slowed down but not blocked in leptotene and zygotene stages. Altogether, the data suggest that *nhr-1* is required for normal meiosis I progression through mechanisms likely involving either the regulation of normal bouquet formation or the maintenance of bouquets at the nuclear envelope.

**Figure 2:**
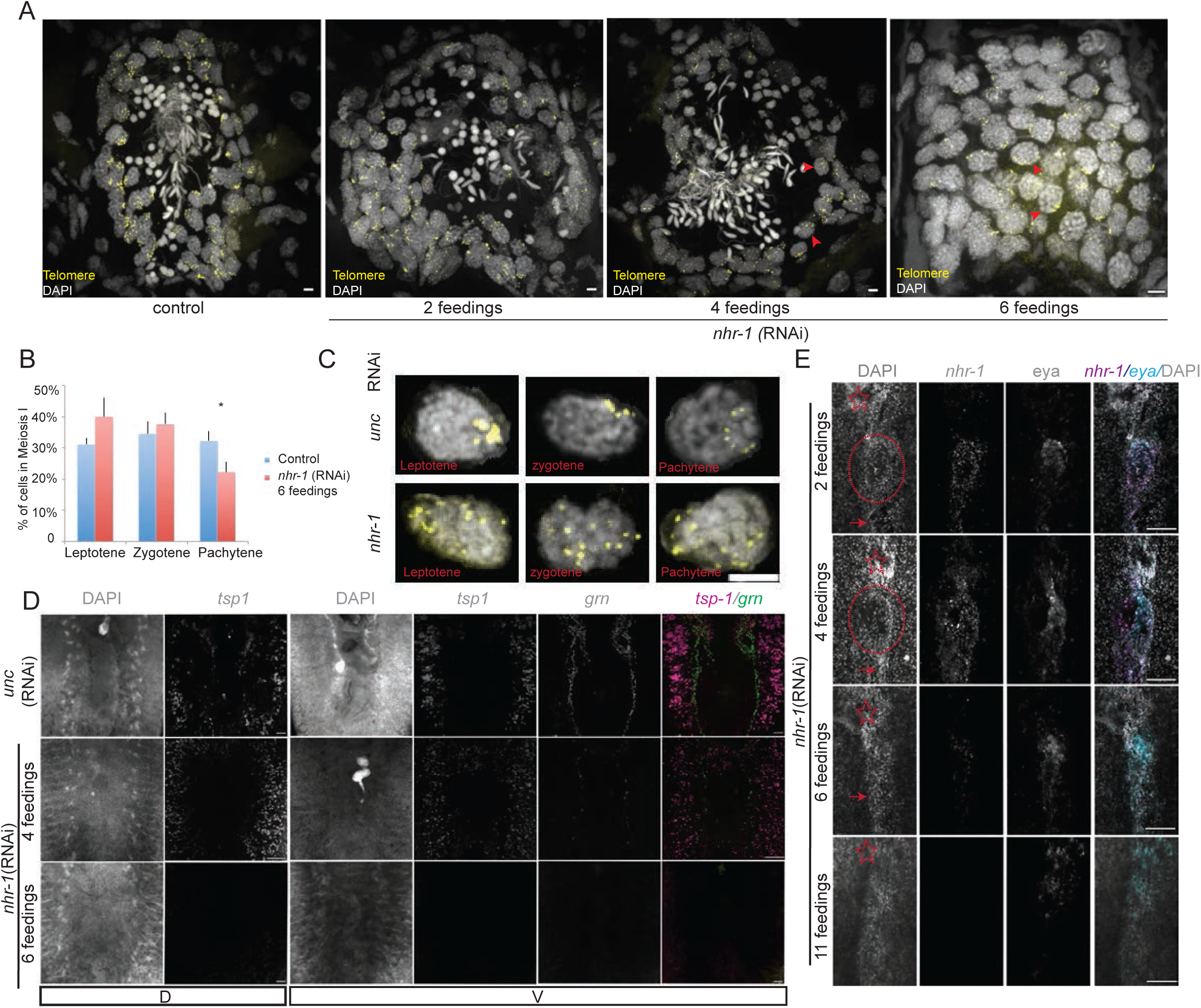
*nhr-1* is necessary for sexual reproductive system maintenance in adult planarians. (A) Nuclear and telomere staining in one nuclear layer in testes. Scale bar:10µm (B) Histogram of leptotene, zygotene and pachytene stage spermatocytes distribution in meiosis I cells observed in half testes by whole mount DAPI staining (n=4 for each sample, *p<0.05). (C) Representative image showing normal bouquet in control animals (upper row) and failed bouquet (lower row) in *nhr-1*(*RNAi*) animals. Scale bar: 5µm (D) Nuclear and FISH staining for gland and sperm duct in control and *nhr-1*(*RNAi*) animal tails. Scale bar: 100µm (E) Nuclear and FISH staining for *nhr-1* expression in oviduct (red arrowheads) in control and *nhr-1*(*RNAi*) animals. Red stars show the brain close to the ovary. Ovaries are in red circles. Scale bar: 20µm

### Expression of known specific marker genes of the RS are affected by *nhr-1*(*RNAi*)

*nhr-1* is expressed in most organs of the adult hermaphroditic planarian RS. In order to gain a better understanding of the dynamics of resorption of the RS after *nhr-1*(*RNAi*), we investigated the expression of known reproductive organ markers during this process. Most *nhr-1*^+^ cells in both male and female gonads expressed *gh4*, a germ stem cell and spermatogonial cell marker (Saberi et al., 2016)(Figure S4B and S4D), but the expression of *nanos* and *nhr-1* did not overlap extensively (Figure S4A and S4C). Given that *nanos* is an evolutionarily conserved marker for germ stem cells (Wang et al., 2007), this result suggested that *nhr-1* is expressed in more differentiated states of the germ stem cells. In fact, after 4 rounds of *RNAi* treatment the ovary appeared vacated of cells, including differentiated oocytes, and by the 6^th^ round of RNAi feeding the ovaries became undetectable (Figure 2E). Co-localization of *nhr-1* with *eyesabsent (eya), grn and tsp-1*(Figure S4E and S4F) showed that *nhr-1* is expressed in oviduct, sperm duct and accessory glands, respectively (Chong et al., 2013; Chong et al., 2011). After 4 rounds of *nhr-1(RNAi), tsp-1* and *grn* showed markedly reduced expression levels while retaining their overall spatial expression patterns, suggesting that *nhr-1* may directly maintain gene expression of *tsp-1* and *grn*. After 6 rounds of RNAi feeding, *tsp-1* and *grn* expression became undetectable (Figure 2D). Interestingly, even though the ovaries became undetectable after the 6^th^ round of RNAi feeding, the oviduct could still be labeled by the expression of *eya* (Figure 2E). Additionally, the expression of *eya* in the brain region persisted even after 11 rounds of *nhr-1 RNAi* feeding (Figure 2E), suggesting that *nhr-1* likely regulates gene expression specifically in the RS. Altogether, our data indicated that *nhr-1* may specifically regulate gene expression in the RS, with gene expression in different RS cells and tissues responding to *nhr-1* loss with different kinetics.

### *nhr-1* is specific and essential for the regeneration of all reproductive system components

To determine whether *nhr-1* may play a role in RS regeneration, we characterized the timing of *nhr-1* expression along with the re-establishment of the accessory reproductive organs in regenerating fragments derived from adult, sexually mature planarians. We followed adult head fragments produced by amputating in the pre-pharyngeal region (Figure 3A, B). These fragments initially retain ovaries and some anterior reproductive organ tissues but are devoid of all RS tissues found in the trunk and tail of the animal after amputation (Figure 3A, B). Head fragments fully regenerate the RS 32 days post amputation (DPA) (Figure 3A). During this process, *nanos*^*+*^ germ stem cells persist, while differentiated gonad cells disappear soon after amputation (Wang et al., 2007). Consistent with data from decapitated fragments (Table S1), detection levels of *nhr-1* expression became sharply reduced after amputation (Figure 3B). However, its expression became noticeable once again 12 DPA, when a newly regenerated pharynx could be detected by DAPI staining, and *nanos*^*+*^ cell had already lined up in a pattern indistinguishable from primordial testes (Figure 3A and 3B). By 12 DPA, *nhr-1* expression was detected in the center of the regenerating tail, where the future dorsal glands will regenerate (Figure 3A and 3B). By 22 DPA, *nhr-1*^*+*^ cells formed a circle in this region (Figure 3A and 3B), and by 28 DPA the marked *nhr-1* expression highlighted the prospective gland structure that eventually became visible by 32 DPA (Figure 3A and 3B). The gland was established after the gonad pore formed, which happened in about half of the tested animals by 32 DPA. It is interesting to note that *nanos*^*+*^ cells started to differentiate 28 DPA, a time when *nhr-1* expression was abundant at the gland region anlagen. These results suggested that *nhr-1* expression precedes the establishment of the definitive RS tissues during regeneration.

**Figure 3:**
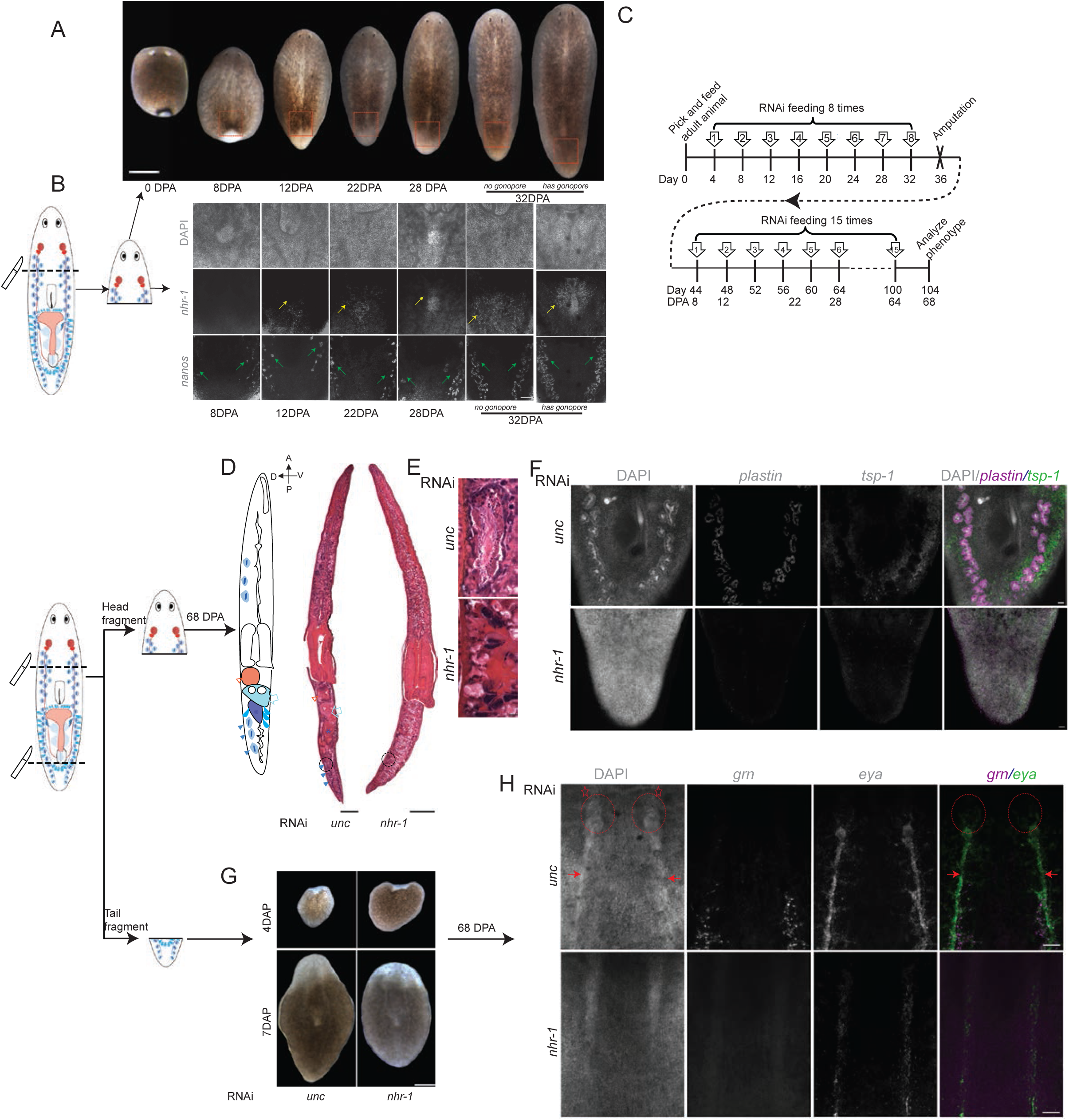
*nhr-1* specifically regulates reproductive system regeneration. (A) Head fragment regeneration time course. Red rectangle indicates the imaging region in panel B. Scale bar: 100µm (B) DAPI and FISH staining for *nhr-1* and *nanos* in the dorsal gland region of the regenerated tail. Yellow arrowhead shows *nhr-1* expression. Green arrowhead shows expression of *nanos*. Scale bar: 100µm (C) Feeding and amputation schedule for testing function of *nhr-1* during regeneration. Adult animals were amputated after 8 rounds of RNAi feeding. 8 days after amputation, the regenerated RNAi animals were fed with RNAi food for 15 more times. (D) Hematoxylin and eosin (H&E) staining of animals regenerated from control *unc-22(RNAi*) and *nhr-1*(*RNAi*) animals 68 days after amputation. Orange triangle shows the copulatory bursa. Blue arrowhead indicates bulbar cavity. Blue asterisk marks penis papilla. Blue triangle points at testis. Scale bar: 100µm (E) Zoom in of the H&E staining in the putative testes region circled in panel D. (F) DAPI and FISH staining for *plastin* and *tsp-1* in dorsal gland region of regenerated tail.Scale bar: 100µm (G) Image of regenerated head from *unc-22(RNAi*) control and *nhr-1*(*RNAi*) tail fragments 4 and 7 days post amputation. Scale bar: 400µm (H) DAPI and FISH staining for *grn* and *eyesabsent (eya*) in the ventral site of regenerated head. Ovaries are in the red circles. Oviducts are pointed by red arrowheads. Red star shows the brain in close proximity to the ovary. Scale bar: 100µm

We proceeded to determine the function of *nhr-1* in RS regeneration by first subjecting sexual adult animals to RNAi for 8 times followed by amputation of heads and tails and further RNAi treatments of these anterior and posterior fragments to chronically sustain *nhr-1* loss of function (Figure 3C, D). Regeneration of the RS in these fragments was examined 68 DPA. In the first week of regeneration, *nhr-1 RNAi* fragments formed blastemas that were indistinguishable from control animals (Figure 3H). While most control animals formed gonopores by 60 DPA, *nhr-1 RNAi* animals had not developed gonopores by 68 DPA (Figure S5A). By 68 DPA, control head fragments had regenerated the full set of mature RS organs, including the copulatory bursa, the bulbar cavity, the penis papilla and testes containing mature spermatids, while the head *nhr-1*(*RNAi*) fragments lacked all of these structures (Figure 3E and 3F). No testis-like structures could be found by DAPI staining on regenerated *nhr-1*(*RNAi*) fragments (Figure 3G), and neither testes nor accessory glands could be detected by the molecular markers *tsp-1* and *plastin* (Figure 3G). We also failed to detect sperm ducts and oviducts by either DAPI or molecular markers on the ventral side (Figure S5C). Nevertheless, gut and pharynx regenerated normally in these fragments with *nhr-1* expression depleted, suggesting the function of this NHR is specific to the RS. Next, we examined the regenerated heads from *nhr-1*(*RNAi*) tail fragments (Figure 3D). Consistently, the *nhr-1*(*RNAi*) fragments failed to regenerate tissues of the RS including the ovary, oviduct and sperm duct (Figure 3I), but regenerated photoreceptors (Figure 3G) as well as the cephalic ganglia of the central nervous system normally (Figure S5B). We conclude from these data that *nhr-1* is specific and essential for the regeneration of the RS in planarians.

### Gene expression profiling of *nhr-1*(*RNAi*) animals revealed both novel genes and metabolic regulators of the RS

To better understand how *nhr-1* maintains tissue homeostasis and regulates regeneration of the RS, we profiled differential gene expression changes between intact sexual adult worms subjected to *nhr-1*(*RNAi*) and *unc-22(RNAi*) controls (Figure 4A). As knocking down of *nhr-1* expression became stronger with increased numbers of RNAi treatments (n=4, 6 and 8), more genes showed reduced expression levels (Figure 4B). The numbers of genes with reduced expression increased from 31 after 4 rounds of feeding to 3,604 after 8 rounds of feeding (Figure 4B). Reported molecular markers of testes, ovaries, and accessory organs were among the genes displaying decreased expression levels after *nhr-1*(*RNAi*) (n=134, Figure 4C), hence validating the quality of our gene set. Expression of molecular markers of germ stem cells (*i.e., nanos, gh4*) were not affected by *nhr-1*(*RNAi*), consistent with previous observation that *nhr-1* is required to maintain differentiated states of germ cells and RS somatic structures (Figures 1 and 2). Gene Ontology analyses of the gene sets with downregulated expression after 6 and 8 rounds of RNAi feeding showed an overrepresentation of functions associated with metabolic processes, especially lipid catabolic process (Figure 4D), including the nutrition sensor AMPK (Chantranupong et al., 2015), Acyl-coenzyme A catabolism-related genes (Acyl-coenzyme A thioesterase, Acyl-CoA-binding domain-containing protein, and Acyl-CoA synthetases) (Tillander et al.,2017), the lipid droplet consuming gene (LPA acyltransferase)(Leung, 2001; Pol et al., 2014), the cholesterol synthesis gene 3-hydroxy-3-methylglutaryl-coenzyme A reductase (HMGCR) (Sharpe and Brown, 2013) and the β-oxidation related genes Acetyl-CoA acetyltransferase, Mitochondrial carnitine/acylcarnitine carrier protein, Peroxisomal carnitine O-octanoyltransferase (Indiveri et al., 2011). Taken together, these results indicate that *nhr-1* likely regulates the tissue homeostasis and regeneration of the RS through the activation of genes associated with discrete metabolic pathways.

**Figure 4:**
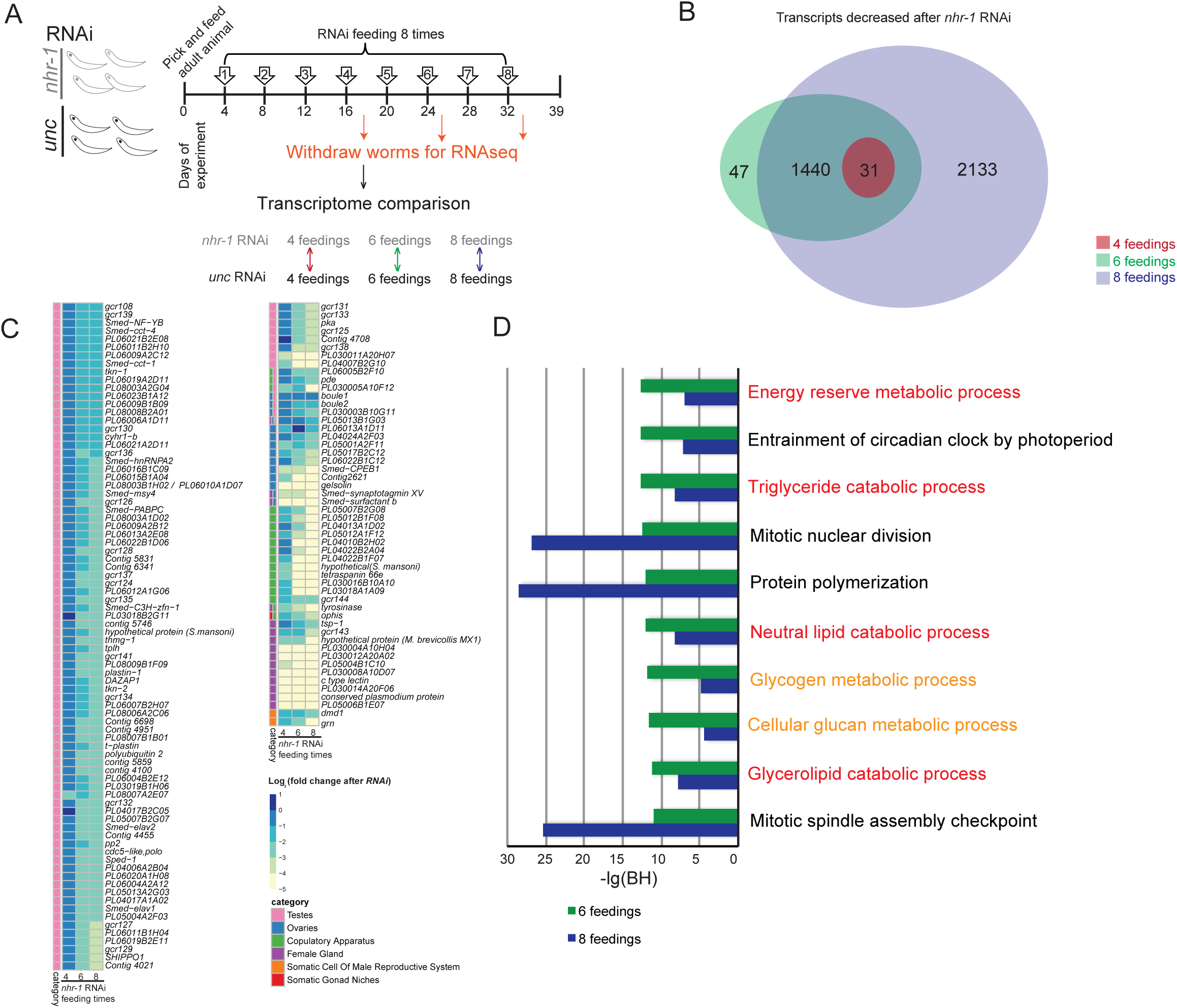
*nhr-1* is necessary for reproductive system and lipid metabolism genes expression. (A) RNAi feeding schedule and sample collection for transcriptome analysis. Both the control (*unc-22* RNAi) and test (*nhr-1* RNAi) animals were collected 2 days after RNAi feeding 4, 6 and 8 times. 2 days after collection, mRNA was extracted from those animals and submitted for sequencing (n=4 replicates, each containing 2-4 planarians). The transcriptome was compared between *unc-22* and *nhr-1* RNAi animals, which were withdrawn at the same time. (B) Venn diagram showing the number of transcripts decreased after *nhr-1* RNAi feedings at different time points (t-test, adj. p<0.05). (C) The fold change of the expression of reproductive system related genes that decreased after *nhr-1* RNAi. (D) The histogram shows the gene ontology (GO) terms with the 10 lowest Benjamini-Hochberg multiple testing correction (BH) value for the transcripts that decreased after *nhr-1* RNAi feeding 6 times. The BH values of those ten terms for the transcripts decreased after *nhr-1* RNAi feeding 8 times are shown in blue. The terms relating to lipid metabolism are shown in red. The terms relating to other metabolic process are shown in orange.

### *nhr-1*(*RNAi*) uncovered novel and specific accessory reproductive organ markers

By profiling gene expression changes in *nhr-1*(*RNAi*) at different times after treatment, we reasoned that genes reduced their expression levels at the earliest time point (*i.e*., 4 times of feeding) could be upstream regulators of different tissues in the RS, or represent tissues that were affected earliest upon loss of *nhr-1* expression. Interestingly, only 31 genes reduced their expression levels after 4 rounds of *nhr-1 RNAi* feedings (Figure 4B), a time when visible changes in the RS were observed in the testes and reproductive glands (Figure 1 and 2). No gene increased at this time point. Among those decreased genes, the expression patterns in the RS were only known for two genes, *tetraspanin 66e* and *PL04022B1F07*(Rouhana et al., 2017). 23 genes were planarian specific, *i.e*., devoid of obvious homologs in other species. 6 genes were homologous to metabolism-related genes, such as Sodium/potassium-transporting ATPase subunit beta-1, Ectonucleoside triphosphate diphosphohydrolase 5 and ACS (Grevengoed et al., 2014; Lingrel et al., 2003; Read et al., 2009). Importantly, all 31 genes have higher expression levels in sexual adult animals, compared to juvenile animals or the asexual line, CIW4. Notably, expression of 30 genes decreased in the first week after amputation (Figure S6C and Table S2). Thus, we hypothesize that this cohort of 31 genes are likely to include specific regulators of different tissue types in the RS.

We successfully cloned the 30 genes identified and examined their expression patterns by whole mount *in situ* hybridization (WISH) (Figure S6A). Consistent with previous observation with PNA staining that the reproductive glands were affected after 4 rounds of *nhr-1* RNAi (Figure 1), we found 23 genes expressed in the posterior reproductive glands. Additionally, 7 genes were expressed in the testes, 4 genes were expressed in the ovaries, 2 genes were expressed in the yolk glands and 1 gene was expressed in the oviduct (Figure S6A). Hence, testes, ovaries, reproductive glands and yolk glands were among those tissues in the RS that were among the earliest affected by loss of *nhr-1* expression. These results reinforce the function that the NHR encoded by *nhr-1* may play a role in activating downstream genes in a cell-autonomous way.

### Novel markers help refine the anatomical description of posterior accessory reproductive glands

Next, we took advantage of the expanded cohort of markers for the reproductive glands uncovered by the transcriptome analyses (n=19) to study the cell type complexity and fine spatial organization of the posterior reproductive glands. We used fluorescent whole mount *in situ* hybridization (FISH) to dissect the expression patterns of the markers and their relative spatial relationships. From single-gene and multi-gene FISH experiments, we found that gland cells have different distribution patterns on dorsal and ventral sides of the tails. Most gland cells are distributed as circles surrounding the copulatory apparatus. Different markers labeled different gland cells which showed distinct distribution domains. We found the expression of the novel gland markers in both dorsal and ventral posterior regions. Based on their expression patterns, we defined dorsal and ventral gland areas and subdivided each of these into 5 spatially-defined regions (Figure 5A and K).

**Figure 5:**
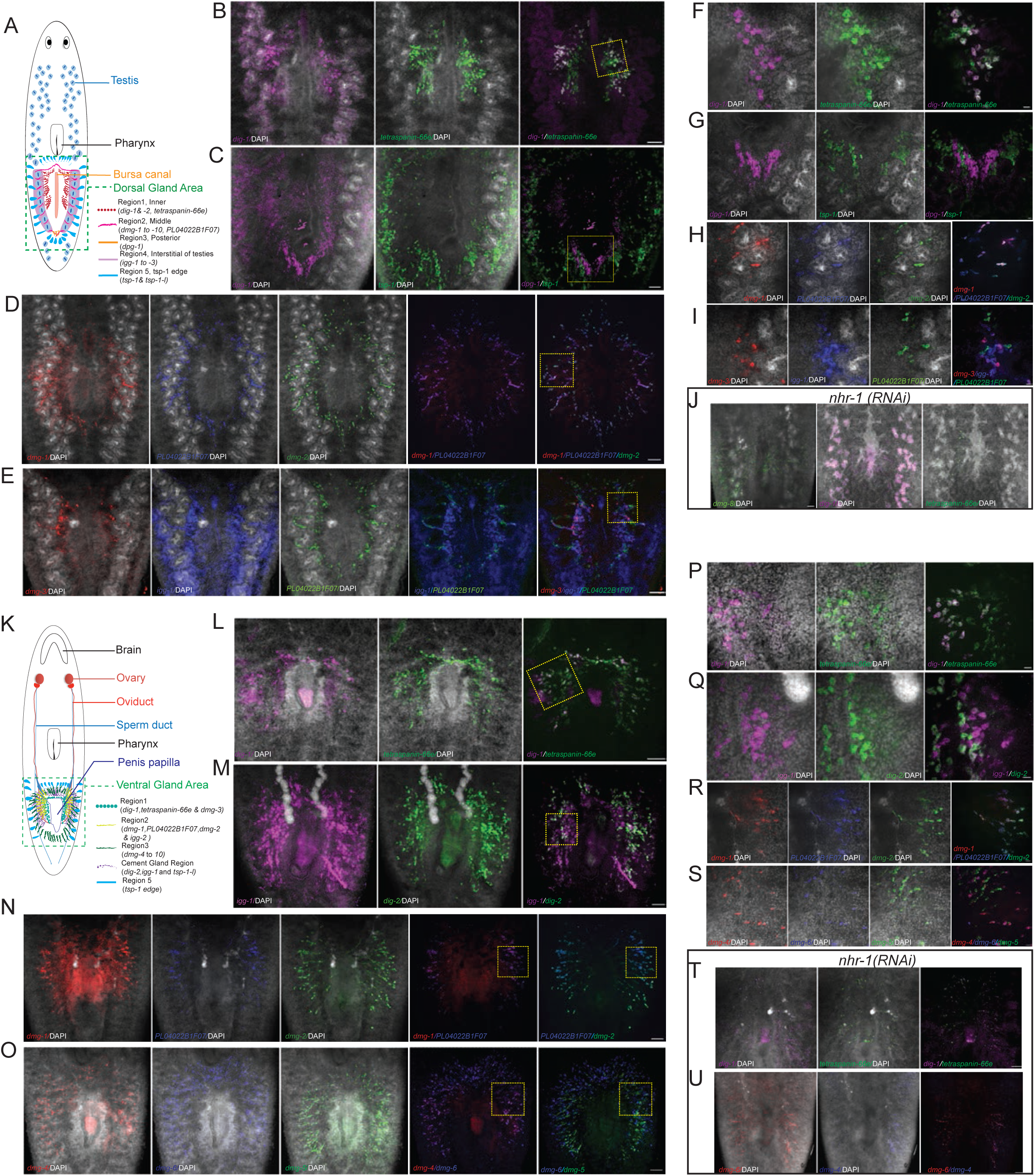
*nhr-1* is essential for novel gland genes expression. (A) The cartoon summarizes the dorsal tail expression pattern of the downstream genes, which decreased after *nhr-1* RNAi feeding 4 times. The glands around the bursa canal can be divided into 5 regions, according to those molecular markers. Examples of genes in each region are shown in panel B to E and further zoomed in single cell layer in panel F to I. (B) FISH for dorsal expression of *dig-1* and *tetraspanin 66e*, which express in region 1. The yellow box shows the region for further zoom in in panel F. Scale bar: 100µm (C) FISH for dorsal expression of *dpg-1*, which expresses in region 3, and *tsp-1*, which express in region 5. The yellow box shows the region for further zoom in in panel G Scale bar: 100µm (D) FISH for dorsal expression of *dmg-1, dmg-2* and *PL04022B1F07*, which express in region 2. The yellow box shows the region for further zoom in in panel H. Scale bar: 100µm (E) FISH for dorsal expression of *dmg-3, igg-1* and *PL04022B1F07*. The yellow box shows the region for further zoom in in panel I. Scale bar: 100µm (F) FISH for dorsal expression of *dig-1* and *tetraspanin 66e* in single cell layer. Scale bar: 20µm (G) FISH for dorsal expression of *dpg-1* and *tsp-1* in single cell layer. Scale bar: 20µm (H) FISH for dorsal expression of *dmg-1, dmg-2* and *PL04022B1F07* in single cell layer. Scale bar: 20µm (I) FISH for dorsal expression of *dmg-3, igg-1* and *PL04022B1F07* in single cell layer. Scale bar: 20µm (J) FISH for expression of *dmg-8, dig-1* and *tetraspanin 66e* in dorsal tail of sexual worm after *nhr-1* RNAi feeding 4 times. Scale bar: 100µm (K) The cartoon summarizes the ventral tail expression pattern of the downstream genes, which decreased after *nhr-1* RNAi feeding 4 times. The glands around the penis papilla can be divided into 5 regions, according to those molecular markers and *tsp-1*. Examples of genes in each region are shown in panel L to O and further zoomed in single cell layer in panel P to S. (L) FISH for ventral expression of *dig-1* and *tetraspanin 66e*, which are in region 1. The yellow box shows the region for further zoom in in panel P. Scale bar: 100µm (M) FISH for ventral expression of *igg-1* and *dig-2*, which express in region 4 or cement gland. The yellow box shows the region for further zoom in in panel Q. Scale bar: 100µm (N) FISH for ventral expression of *dmg-1, PL04022B1F07* and *dmg-2*, which express in region 2. The yellow box shows the region for further zoom in in panel R. Scale bar: 100µm (O) FISH for ventral expression of *dmg-4, dmg-5* and *dmg-6*, which express in region 3. The yellow box shows the region for further zoom in in panel S. Scale bar: 100µm (P) FISH for ventral expression of *dig-1* and *tetraspanin 66e* in single cell layer. Scale bar: 20µm (Q) FISH for ventral expression of *igg-1* and *dig-2* in single cell layer. Scale bar: 20µm (R) FISH for ventral expression of *dmg-1, PL04022B1F07* and *dmg-2* in single cell layer. Scale bar: 20µm (S) FISH for ventral expression of *dmg-4, dmg-5* and *dmg-6* in single cell layer. Scale bar: 20µm (T) FISH for expression of *dig-1* and *tetraspanin 66e* in ventral tail of sexual worm after *nhr-1* RNAi feeding 4 times. Scale bar: 100µm (U) FISH for expression of *dmg-4* and *dmg-6* in ventral tail of sexual worm after *nhr-1* RNAi feeding 4 times. Scale bar: 100µm

The 19 gland markers subdivided the dorsal gland area into 5 discrete regions which allowed us to name the novel genes after their detected spatial location. Region 1 was defined by the *dorsal inner gland 1 and 2* genes (*dig-1 and dig-2*) and *tetraspanin-66e*, which were expressed in the dorsal regions lateral to the bursa canal (Figure 5B and Figure S7A). *dig-1* co-localized with *tetraspanin-66e* in the exterior part of the *tetraspanin-66e* ^*+*^ domain (Figure 5F), while *dig-2* co-localized with *tetraspanin-66e* in the anterior part of the *tetraspanin-66e*^+^ domain (Figure S7A). Region 2 was defined by the *dorsal middle gland* genes 1 through 10 (*dmg-1* to *dmg-10*) and *PL04022B1F07*, all of which were expressed in more exterior circles relative to *dig-1 and -2* and *tetraspanin-66e*. Gland cells in Region 2 marked by these genes were distributed around the testes, dorsal and ventral to the bursa canal (Figure 5A). *dmg-1* and *PL04022B1F07* were co-expressed in the same gland cells, while *dmg-2* was expressed in a broader domain (Figure 5D and 5H). *dmg-3* was expressed in a smaller, distinct domain that does not overlap with *PL04022B1F07* expression (Figure 5E an 5I). *dmg-4* and *dmg-5* were expressed in the same cells, which were different from those cells expressing *dmg-6* (FigureS7B and S7F). *dmg-7* and *dmg-8* were co-expressed in the same cells (Figure S7C and S7G). Region 3 was defined by the *dorsal posterior gland 1* gene (*dpg-1*), which was expressed in a small number of gland cells posterior to the bursa canal (Figure 5A, 5C, 5G and FigureS6A). Region 4 was determined by the *interstitial gland* genes (*igg-1, -2* and -*3*), which were robustly expressed in a very broad domain wrapping around the posterior testes (Figure 5A). *igg-1* showed partial colocalization with *dmg-3*, but not *PL04022B1F07* (Figure 5E and 5I), while *igg-2* labeled the more exterior part of the *igg-3* domain (FigureS7D and S7H). Region 5 was specified by one gene which we named *tsp-1-like* (*i.e., tsp-1-l*) as it is expressed in similar domains as *tsp-1* (Figure S6A). Interestingly, neither *dpg-1*, nor *igg-1* overlapped with *tsp-1*, which was widely expressed in the dorsal reproductive gland (Figure 5C and 5G, Figure S7E).

The anatomical distribution of the reproductive glands in the ventral side of the worm could likewise be divided into 5 regions according to the spatial expression pattern of the above described genes. Ventral Region 1 was defined by *dig-1, tetraspanin-66e* and *dmg-3* which were expressed in the gland next to the penis papilla, including 1-2 layers of cells located in front of this gland (Figure 5K, L and S7K), with *dig-1* expressed in the exterior part of *tetraspanin-66e* ^*+*^ cells (Figure 5P). Ventral Region 2 was characterized by the expression patterns of *dmg-1, PL04022B1F07, dmg-2* and *igg-2* in the gland close to the end of the seminal vesicles (Figure 5N and S7I). Similar to their expression in the dorsal side, *dmg-1* and *PL04022B1F07* were expressed in the same gland cells, while *dmg-2* was expressed in more peripheral cells (Figure 5N and 5R). Ventral Region 3 was defined by 7 transcripts expressed around the copulatory apparatus (Figure 5K, O and S7J). Unlike the dorsal side, *dmg-4* and *dmg-6* were likely expressed in the same gland cells, while *dmg-5* was expressed in different cells (Figure 5O, S). *dmg-7* and *dmg-8* were likely expressed in the same gland cells (S7J, M). The fourth ventral or cement gland region was defined by *dig-2, igg-1* and *tsp-1-l* expression (Figure 5K and 5M). We found that the majority of the *dig-2*^*+*^ cells did not express *igg-1* (Figure 5Q), and that *igg-1* was not expressed in *tsp-1*^*+*^ cells (Figure S7L). *igg-3 and dpg-1* were not detected in the ventral region (Figure S7I for *ig3*).

Consistent with the transcriptome analyses (Figure 4B), expression in adult animals of these 30 transcripts decreased after four *nhr-1*(*RNAi*) treatments (Fig 5J, T, U and S8). The expression of *dig-1* in the gland region decreased, while its expression on the testes did not show appreciable changes (Fig 5J, T). This suggested that *dig-1* expression in the gland is likely to be specifically regulated by *nhr-1*. *dig-11, dig-2, tetraspanin-66e, dmg-6, dmg-8, igg-2* and *igg-3* also changed their expression patterns after *nhr-1*(*RNAi*) (Figures 5J, T, U, and S8A, E, F and C). In the dorsal regions, expression of both *tetraspanin-66e* and *dig-2* extended to the dorsal middle region (Fig S8A), while *dmg-8, igg-2* and *igg-3* expanded their expression to the inner region (Figure 5J and S8C). In the ventral side, *dig-1* and *tetraspanin-66e* extend to the region around the end of seminal vesicles (Fig 5T), while the expression of *dmg-6* and *dmg-8* were now detected in the central region (Figure 5U and S8F). These data suggest that *nhr-1* is not only essential for the development and regeneration of the RS, but also for regulating anatomical expression domains of the accessory reproductive organs.

### *Smed-acs-1* is essential for development, maintenance and regeneration of the RS

To determine the functions of the 30 genes responding earliest to loss of *nhr-1* expression, we carried out RNAi screening in sexual adults and examined for defects in the RS. Of these, we focused on the planarian homolog of Acyl-CoA synthetase *Smed-acs-1,* as we were struck to find out that its silencing by RNAi phenocopied the defects observed for *nhr-1*(*RNAi*). PNA staining showed that both dorsal and ventral glands degenerated after 6 rounds of *acs-1(RNAi*) treatments. After 9 rounds of RNAi, the glands became undetectable (Figure 6A). Degeneration or loss of other tissues in the RS, including the penis papilla, testes, ovaries and oviduct were also evident (Figure 6B). Moreover, the treated worms stopped laying egg capsules after *acs-1(RNAi*) treatments.

**Figure 6:**
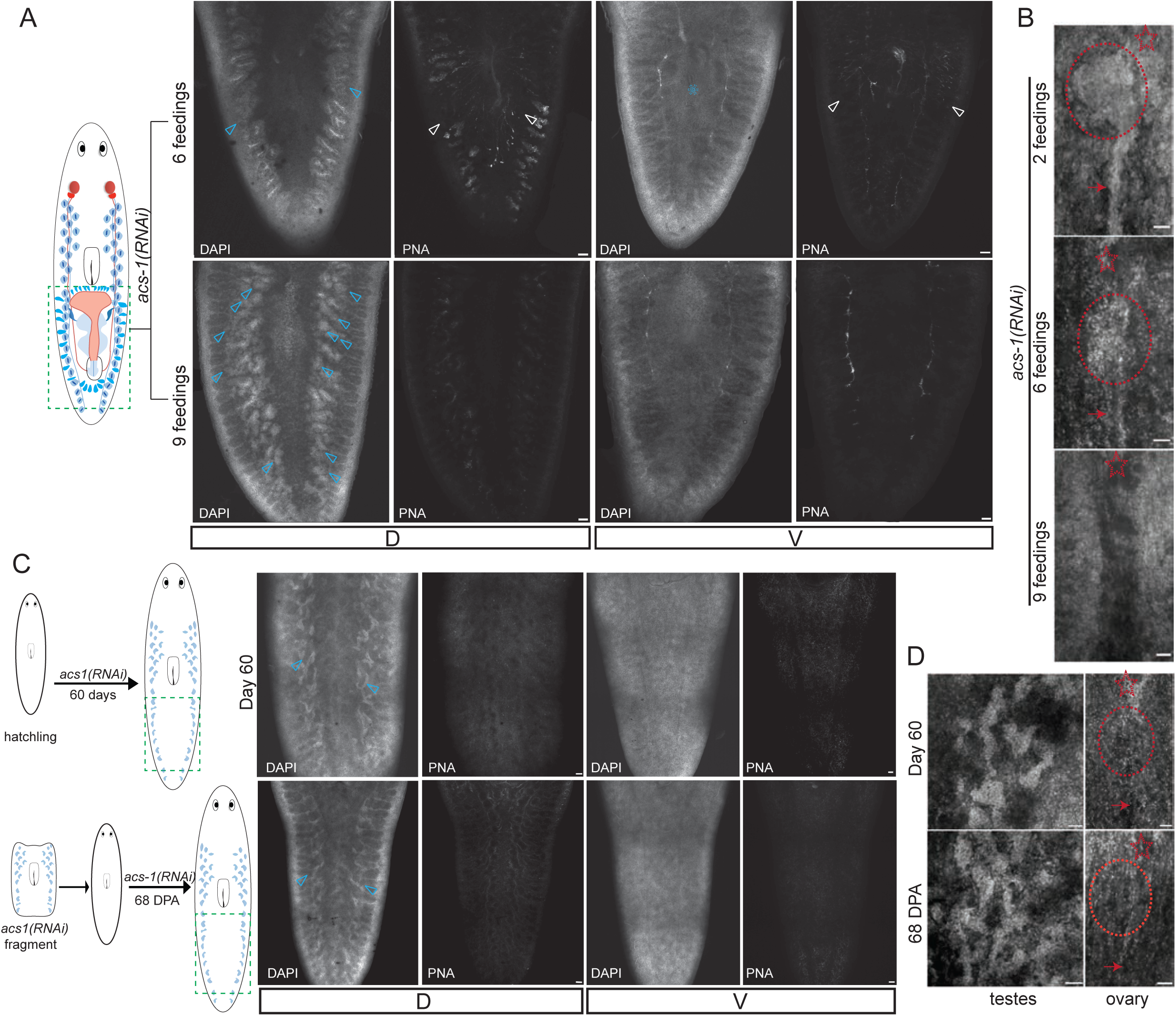
*Smed-acs-1*, a downstream gene of *nhr-1*, is essential for reproductive system development, maintenance and regeneration. (A)DAPI and PNA staining of the tail region in sexual mature planarian after *Smed-acs-1* RNAi feeding 6 and 9 times. The blue triangles point to the testes without mature spermatozoa. The white triangles point to the regressing gland stained by PNA. The blue snow flake marks the penis papilla. Scale bar: 100µm (B)DAPI staining of the ovary and oviduct region in sexual mature planarian after *Smed-acs-1* RNAi feeding 2, 6 and 9 times. The red circle marks the ovary. The red arrow head points to the oviduct. Scale bar: 50µm (C)DAPI and PNA staining of the tail region in grow-up hatchlings (above row) and regenerated worms (below row) after *Smed-acs-1* RNAi feeding. The blue triangles point to the testes without mature spermatozoa. Scale bar: 100µm (D)DAPI staining of the testes and ovaries region in grow-up hatchlings (above row) and regenerated worms (below row) after *Smed-acs-1* RNAi feeding. The red circle marks the ovary. The red arrow head points to the oviduct. Scale bar: 50µm

*acs-1* also shared similar expression dynamics and functions with *nhr-1* during development and regeneration of the RS. *acs-1* expression increased during sexual maturation and decreased after amputation (Figure S9A). Remarkably, body growth and somatic tissue regeneration were not influenced by *Smed-acs-1(RNAi*) in hatchlings and regeneration fragments; however, the reproductive accessory glands were not detectable in both conditions (Figure 6C). The penis papilla and gonopore neither developed, nor regenerated in the *Smed-acs-1(RNAi*) animals (Figure 6C). Similarly, neither the ovaries nor the oviduct developed or regenerated (Figure 6D). Interestingly, the testes both developed and were regenerated after amputation, but failed to produce mature spermatozoa (Figure 6C and 6D). We conclude from these results that *acs-1,* like *nhr-1*, is essential for both RS development and regeneration.

The identified planarian *acs-1* is a homolog of ACS, which is known to be important in fatty acid metabolism (Ellis et al., 2010). *acs-1* codes for a protein containing an AMP-forming/AMP-acid ligase II domain at its N-terminus and phosphopantetheine attachment sites at its C-terminus. The deduced Smed-ACS-1 protein sequence in sexual and asexual planarian share 99.0% identity, yet when compared to sexual animals, the expression of *Smed-acs-1* is higher in sexual than in asexual planarians (Figure S9A). Unlike *nhr-1, acs-1* expression is restricted to comparatively smaller RS domains. *acs-1* is expressed around the testes region of the planarian dorsal side (Figure S9B and S9C) and in the ovary and yolk gland cells on the ventral side (Figure S9B and S9D). These data indicate that *acs-1* function has been restricted to sexual organs and suggest that *acs-1* may regulate RS functions non-cell autonomously.

### *nhr-1* regulates reproductive system maintenance through lipid metabolism

Given that genes regulating lipid metabolism decreased their expression after *nhr-1*(*RNAi*), and that ACS is important for fatty acid uptake, we tested whether *nhr-1* or *acs-1* regulated the RS by modulating lipid metabbolism. We first assayed dietary lipid accumulation in sexually mature planarians using Oil Red O staining. Lipid droplets were quantified 7 days after feeding animals with liver (Material and Methods). Well-fed wild type and *unc-22(RNAi*) animals rarely, if at all, accumulated detectable amounts of lipids (Figure 7A, B), suggesting that under normal conditions planarians efficiently uptake and metabolize dietary lipids. In contrast, animals treated with either repeated rounds of *nhr-1*(*RNAi*) or *acs-1(RNAi*) displayed readily detectable lipid accumulation in the form of lipid droplets (Figure 7A, C), with the most notable accumulation being observed in the areas around the intestine and testes (Figure 7B, D). Interestingly, neither *nhr-1(RNAi),* nor *acs-1(RNAi*), had measurable effects on lipid accumulation in asexual worms (Figure S10A). Thus, we conclude from these data that both *nhr-1* and *Smed-acs-1* are essential for lipid accumulation specifically in sexual adult planarians.

**Figure 7:**
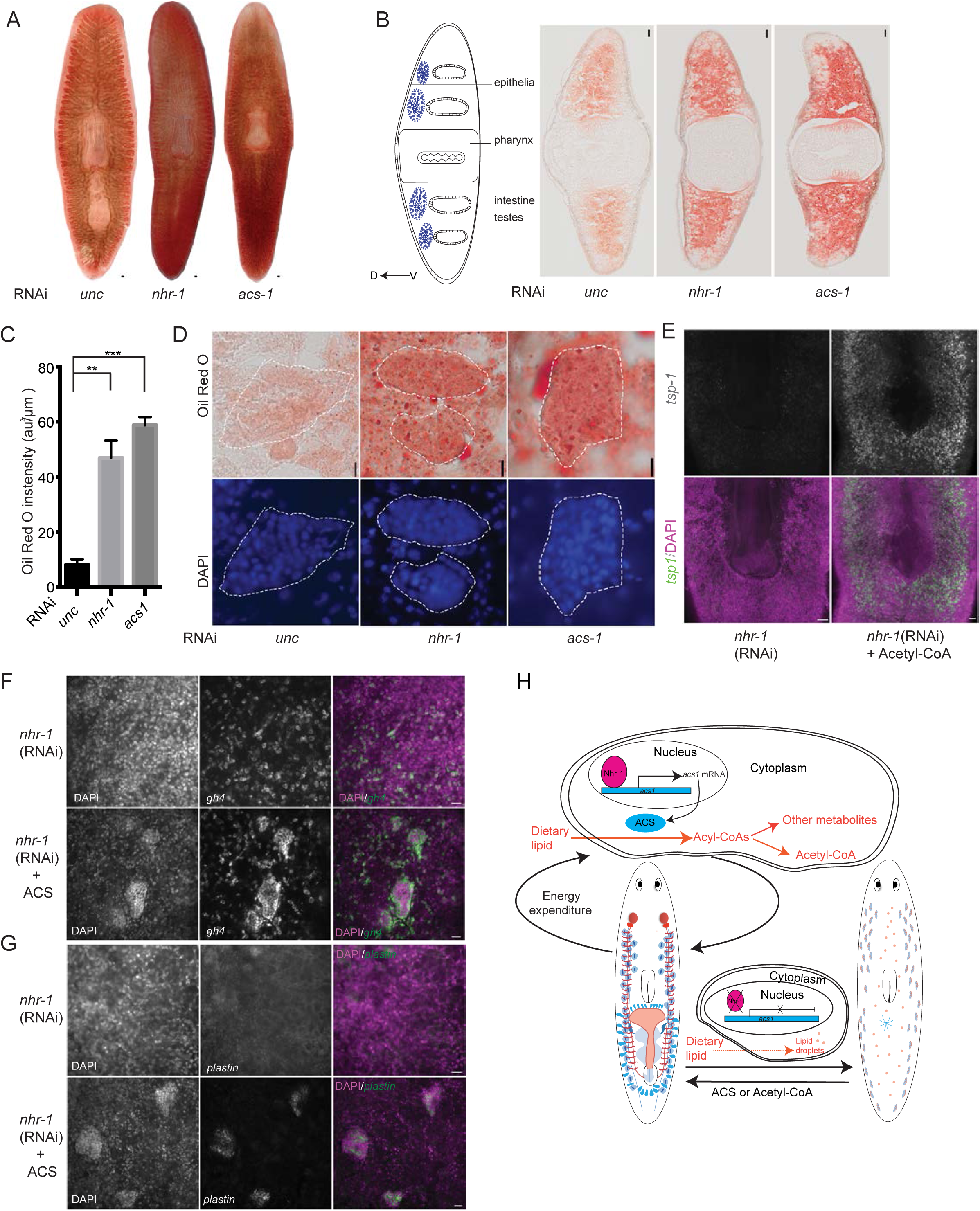
*nhr-1* maintains planarian reproductive system through regulating lipid metabolism (A)Oil Red O staining of the sexual adult planarian 7 days after *unc-22, nhr-*1 and *Smed-acs-1* RNAi feeding. Scale bar: 100µm (B)Oil Red O staining on the transverse sections of the sexual adult planarian 7 days after RNAi feeding. The left illustration shows the structure of the transverse section. Scale bar: 50µm (C)Histogram shows the Mean±SEM value of the Oil Red O staining intensity in the tails (n=3, t-test **p<0.01, ***p<0.001). (D)Oil Red O and DAPI staining around testes region. White dash line circles the testis region according to DAPI staining. Scale bar: 10µm (E)WISH of *tsp-1* and DAPI staining of the dorsal tail region in sexual planarian after long term *nhr-*1 *RNAi* feeding with (right) and without (left) Acetyl-CoA rescue. Scale bar: 100µm (F)WISH of *gh4* and DAPI staining of the testes region in sexual planarian after long term *nhr-*1 RNAi feeding with (below) and without (above) Acyl-CoA synthetase (ACS) rescue. Scale bar: 20µm (G)WISH of *plastin* and DAPI staining of the testes region in sexual planarian after long term *nhr-*1 RNAi feeding with (below) and without (above) Acyl-CoA synthetase (ACS) rescue. Scale bar: 20µm (H)Proposed model for reproductive system maintenance in sexual mature animals via am *nhr-1* and lipid metabolism axis.

Because Acyl-CoA Synthetase is a critical enzyme for fatty acid activation (Ellis et al., 2010), we attempted to rescue the RS by restoring Acyl-CoA Synthetase or lipid metabolites in the sexual adults fed with *nhr-1*(*RNAi*) food. Fatty acid can be converted by the Acyl-CoA Synthetase *in vitro* (Knoll et al., 1994). Planarian RNAi food is a homogenate of beef liver and bacteria, which together with free fatty acids from planarian tissues may offer the substrates for fatty acid conversion. After feeding adult worms with *nhr-1*(*RNAi*) food for 4 weeks, the Acyl-CoA Synthetase from *Pseudomonas* was added to the RNAi feeding schedule for the remaining 6 weeks of the experiment. After 10 weeks of *nhr-1*(*RNAi*) treatment without Acyl-CoA Synthetase supplementation, adult animals lost their testes and accessory reproductive glands as before (7 out of 8) (Figure 7E and S10B). Remarkably, the animals supplemented with Acyl-CoA Synthetase retained their testes (Figure S10B) and gland cells (Figure S10D) indicating that addition of this enzyme was sufficient to rescue the loss of the RS in *nhr-1*(*RNAi*)-treated animals (n=7 out of 7). Additionally, after long-term *nhr-1*(*RNAi*) feedings, clusters of germ stem cells, spermatogonia and other meiotic cells were not detectable by either *gh4* or *plastin* staining (Figure 7F and 7G). However, clusters of *gh4*^*+*^ and *plastin*^*+*^ cells were restored after *nhr-1*(*RNAi*) was provided (Figure 7F and 7G), which suggested that Acyl-CoA Synthetase rescued both the proliferation and differentiation of germ stem cells in the absence of *nhr-1* function. To further evaluate this discovery, we asked if the *nhr-1*(*RNAi*) RS phenotypes could be rescued by the exogenous addition of the metabolite Acetyl-CoA, which is made by Acyl-CoA Synthetase and has been shown to be taken up from the extracellular space by transporters (Pietrocola et al., 2015). We fed or injected Acetyl-CoA to *nhr-1*(*RNAi*) animals and the results were consistent with Acyl-CoA Synthetase feeding, with 5 out of 8 animals displaying rescue of the phenotype as illustrated by the recovery of *tsp-1 expression* in dorsal gland cells and of *plastin* and *gh4* cells in the dorsal testes region (Figure 7E, S10B and S10C). Taken together, increasing an enzyme or exogenously providing a metabolite essential for lipid metabolism was sufficient in both cases to rescue the RS defects caused by *nhr-1(RNAi*) in planarians. These results demonstrated that *nhr-1* likely regulates RS maintenance and regeneration via an Acyl-CoA Synthetase-activated lipid uptake process.

## DISCUSSION

We have shown that both male and female RS components of the planarian *S. mediterranea* are actively maintained by the nuclear hormone receptor *nhr-1* and its downstream target *acs-1* through lipid metabolic pathways (Figure 7H). In many organisms, including humans, the energetic costs associated with reproduction are considerable and rely in great measure on fat metabolism, the major source of energy in animals (Bronson, 1989; Valencak et al., 2009; Wang et al., 2008). In planarians, *nhr-1* activates lipid metabolic genes in sexually mature animals (Figure 4D), including a gene homologous to Acyl-CoA Synthetase (*Smed-acs-1*). During the reproduction of many animals, fat reserves are mobilized, but reducing or abolishing reproduction increases lipid storage in many species (Corona et al., 2009; Judd et al., 2011). Dietary lipids are converted to Acyl-CoAs by Acyl-CoA Synthetase, a role likely played by *Smed-acs-1* in the RS of planarians, where their β-oxidation yields Acetyl-CoA to fuel both reproduction and the lifelong maintenance of the RS. Without *nhr-1*, the expression of *acs-1* is inhibited. Free lipids fail to be taken up by the RS and accumulate in the body, followed by dramatic hypogonadism and the general resorption of all accessory reproductive glands in planarians (Figure 7H). Our study, therefore, has uncovered a reproductive-endocrine signaling axis causally linked to dietary lipid metabolism.

### Lipid metabolism plays a critical role in RS maintenance and regeneration

The insulin and neuroendocrine pathways have been previously shown to be required for sexual maturation and germ cell differentiation in planarians. In *S. mediterranea*, the insulin pathway is activated by insulin like pheromone (*ilp-1*) through its receptor (*inr-1*), and influences both body size and reproductive organs (Miller and Newmark, 2012). The neuroendocrine pathway involves the neuropeptide NPY-8 and its cognate G protein-coupled receptor NPYR-1, and their knockdown results in the loss of differentiated germ cells and sexual maturity (Saberi et al., 2016). Although the insulin and lipid metabolism pathways are known to crosstalk in other species, the lipid metabolic processes regulated by *nhr-1* are likely independent of the insulin pathway. Even though *ilp-1* expression decreased after 6 rounds of *nhr-1*(*RNAi*) treatment, neither *nhr-1*, nor *Smed-acs-1*, influenced body size even after 12 feedings. Similarly, *nhr-1* does not respond to NPY8, suggesting that *nhr-1* is likely not part of the neuropeptide pathway, an observation supported by the fact that NPYR-1 is the receptor of NPY-8. Moreover, neither the insulin, nor the neuropeptide pathway-related genes altered their expression after *nhr-1*(*RNAi*). These data suggest that the lipid metabolism regulated by *nhr-1* is a third and novel pathway that plays a critical role in the reproductive system of *S. mediterranea*.

*nhr-1,* its putative target *acs-1* and the downstream lipid metabolic genes and metabolites may also have conserved roles in other species of the Lophotrochozoa. In planarian *Dugesia ryukyuensis*, yolk gland regeneration is inhibited by excess 17β-estradiol (Miyashita et al., 2011), which is a lipid catabolite (Wollam and Antebi, 2011) and a key regulator of lipid metabolism (Sieber and Spradling, 2015). Also, the hydrophobic fraction of tissue homogenates from the planarian *Polycelis nigra* is known to induce the sexual state in the asexual *Dugesia gonocephala sensu lato* (Grasso et al., 1975), which suggests that lipid and its metabolites are important for the sexual maturation and reproduction. In *Schistosoma mansoni*, fatty acid oxidation and acyl-CoA synthetase are required for egg production (Huang et al., 2012). Both *nhr-1* and *Smed-acs-1* have conserved homologs in other free-living and parasitic platyhelminthes. Our work suggests that antagonists to *nhr-1* and/or *Smed-acs-1* may repress sexual reproduction in the Lophotochozoa by desexualization and thus may be useful drug targets for parasitic control.

### *nhr-1* is required for lipid metabolism in the planarian reproductive organs

*nhr-1* likely directs the expression of multiple downstream genes. The protein encoded by *nhr-1* has two conserved nuclear hormone receptor DNA binding domains in the N terminal and one ligand binding domain in the C terminal. After *nhr-1*(*RNAi*), the majority of affected genes showed significantly downregulated expression suggesting that *nhr-1* mostly activates gene transcription. In fact, the expression of only 31 genes appeared to be significantly affected when defects were first observed during the early stages of *nhr-1*(*RNAi*) treatment. Most of these genes were predominantly expressed in the dorsal and ventral glands around the copulatory organs (Figure 5). *nhr-1* is a lophotrochozoan-specific nuclear receptor (Tharp et al., 2014), but a homologue to the retinoid-related orphan receptor β (RORβ). Members of the ROR family of receptors such as RORγt can be activated by oxysterols, which are cholesterol derivatives produced during lipid metabolism (Soroosh et al., 2014). Given the low levels of *nhr-*1 and *acs-1* expression in the asexually reproducing planarians, and that glands are important organs for sterol synthesis (Kurzchalia and Ward, 2003; Niwa and Niwa, 2016; Wollam and Antebi, 2011), we speculate that a likely source of the unknown ligand for *nhr-1* may originate from the accessory gland organs of the RS. Because *nhr-1* expression is dramatically decreased after amputation and reactivated during RS regeneration a future comparison of the metabolome between these two stages may help identify the ligands for *nhr-1*.

### A reproductive-endocrine signaling axis is required for the maintenance and regeneration of the RS

Our data indicate that *nhr-1* triggers conserved lipid metabolism pathways to maintain the planarian RS. Depletion of *nhr-1* expression during homeostasis decreased the expression levels of lipopolysaccharide and other downstream putative target genes, such as tumour necrosis factor (TNF)α and interleukin (IL), which are known to initiate liver regeneration after partial hepatectomy(Taub, 2004). Additionally, our studies revealed a downregulation of *caveolin* and *HMGCR* gene expression after *nhr-1*(*RNAi*), both of which are also involved in lipid metabolism and are essential for liver regeneration (Delgado-Coello et al., 2011; Fernandez et al., 2006; Gazit et al., 2010; Trentalance et al., 1984). Additionally, recent work has showed that intestine regeneration is also be boosted by dietary lipid(Beyaz et al., 2016). Thus, the molecular pathway of RS regeneration, which is triggered by *nhr-1* and regulated by *Smed-acs-1*, may function in different organs. Altogether, we take these data to suggest that the NHR-dependent regeneration of the RS may not be a unique feature of this system and that other organ-endocrine axes may exist that are mediated by other not yet characterized NHRs encoded by the *S. mediterranea* genome.

### *Smed-acs-1* specifically regulates lipid metabolism in RS

ACS can convert fatty acids to different products, which may have different metabolic fates in different tissues (Grevengoed et al., 2014). In vertebrates, long ACS isoform 4 (ACSL4), fatty acid transport protein 3 (FATP3), ACS bubblegum 1 (ACSBg1) and ACS bubblegum 2 (ACSBg2) are enriched in the RS, producing different types of Acyl-CoAs (Grevengoed et al., 2014). Though defects in brain were observed in ACSL4 and ACSBg mutants, defects in the RS were not reported in animals lacking a single ACS (Grevengoed et al., 2014), suggesting that different organs may depend on specific metabolites for their normal function. Given the very specific expression patterns of *acs-1* in the adult planarian, it is likely that *acs-1* may be non-cell autonomously regulating the maintenance and regeneration of the planarian RS. Based on gene expression, the most likely tissues endowed with high *ACS-1* activity are the ends of gut branches, the exterior of the lobes of testes, yolk glands and glands surrounding the ovaries. These cell types may convert dietary lipids from the intestine and transport the Acyl-CoAs to other tissues of the RS. That such transport of metabolites likely exists in planarians is supported by the rescue experiments in which we fed either bacterial ACS or Acetyl-CoA to *nhr-1*(*RNAi*)-treated animals (Figure 7). In fact, the rescue of the defects in the RS in *nhr-1*(*RNAi*) planarians by exogenously administered bacterial ACS, suggests that planarian RS may not necessarily rely on a single kind of Acyl-CoAs.

Because ACS protein and Acetyl-CoA could rescue defects in the RS after *nhr-1*(*RNAi*), we suspect that the AMPK and Acetyl-CoA synthesis pathways in both mitochondria and cytosol may be controlled by *acs-1* in the planarian RS. After fatty acid is converted to Acyl-CoAs by ACS in the cytosol, Acyl-CoAs can be transported by mitochondrial carnitine/acylcarnitine carrier protein (SLC25A20) to synthesize Acetyl-CoA to be used by the TCA cycle to generate ATP. Alternatively, Acyl-CoAs can be used directly in the cytosol to produce Acetyl-CoA, which is further used to generate sterols by HMGCR (Pietrocola et al., 2015). After 6 to 8 feedings of *nhr-1*(*RNAi*), the expression of key molecules involved in Acetyl-CoA generation and consumption in both mitochondria and cytosol decreased significantly (Figure 4). AMPK expression also decreased after 6 rounds of *nhr-1*(*RNAi*). AMPK can be activated by fatty acid and modulate meiosis in the gonads of *Drosophila, C. elegans* and mice (Bertoldo et al., 2015; Ellis et al., 2010). Given that and *acs-1(RNAi*) phenocopies the RS defects of *nhr-1*(*RNAi*), it is likely that AMPK may be a downstream target of *acs-1* to regulate planarian gonad functions.

### Implications for reproduction, fat metabolism and lifespan

In many organisms, increased life span is associated with reduced reproduction, but drastically increased lipid storage and generally improved survival under starvation conditions (Rion and Kawecki, 2007; Tatar et al., 2001). As such, it has been postulated that lipids set aside for reproduction are unavailable to support the maintenance and survival of other somatic tissues, ultimately diminishing the life-span of the organism (Shanley and Kirkwood, 2000). This tradeoff between reproduction and the homeostasis of other somatic functions has led to the notion that individuals with curtailed reproduction survive better and live longer than those with higher reproductive output and vice versa (Partridge et al., 2005; Reznick et al., 2000). Yet, how reproduction, fat metabolism, and life span may or may not be mechanistically intertwined remains poorly understood (Hansen et al., 2013). The findings reported here indicate that planarians may provide a novel biological context in which to unravel this issue.

Unlike the most actively researched organisms used to study reproduction, lipid metabolism and aging (*i.e*., flies, nematodes and vertebrates), planarians are negligibly senescent and can lose and regenerate their entire RS seemingly endlessly. When either starved or amputated, the hermaphroditic sexual organs are resorbed likely due to loss of *nhr-1* signaling, only to be restored once food is available, the animals reach an appropriate size (Newmark and Sanchez Alvarado, 2002), and lipid metabolism is once again activated to maintain the RS. In both men and women, the reduction of hormone synthesis and gametes are usually the earliest signs of aging (Gunes et al., 2016; Makabe et al., 1998), with other organs such as the liver and brain decreasing in volume in the elderly (Hedman et al., 2012; Makabe et al., 1998; Motta et al., 2002; Paniagua et al., 1991; Tajiri and Shimizu, 2013). Moreover, the ectopic lipid accumulation observed in normal aged people and premature menopause patients (Knauff et al., 2008; Toth and Tchernof, 2000), is only observed in planarians after the loss of either *nhr-1* or *acs-1*. Thus, decline of fat oxidation and lipid synthesis may be a general cause of aging and resorption of organs. The activation of *nhr-1* orthologs, ACS, or other downstream associated genes that may rebalance lipid metabolism may result in the rejuvenation of organs. As such, planarians present a unique opportunity to both determine whether steroid hormones play a critical role in the interconnection between reproduction, lifespan and fat metabolism, and to help establish how these three processes may be causally connected.

## ACKNOWLEDGEMENTS

We thank E. Reid and F. Walker for help in cloning and screening, all members of the A.S.A. laboratory for discussion and advice. We also thank Dr. Y. Wang, N. Thomas and other histologists in the Stowers Institute Histology Facility for their beautiful sectioning and staining work. We acknowledge S. M. Merryman and other members at the Stowers Institute Planarian Core Facility for planarian husbandry. We also acknowledge Zulin Yu, Jeffrey Lange and other members at the Stowers Institute Microscopy Facility for assistance. A.S.A. is a Howard Hughes Medical Institute and Stowers Institute for Medical Research investigator. This work was supported in part by NIH grant R37GM057260.

## Experimental Procedures

### Planarian culture and RNAi feeding

Sexual *S. mediterranea* were maintained at 20°C as previously described(Guo, Zhang, Rubinstein, Ross, & Alvarado, 2016). Worms with gonopore and about 10mm length were used as sexually mature animal for experiments. The juvenile worms were ∼4mm length without gonopore.

### RNA extraction and gene expression analyses

RNA of the *nhr-1* and *unc* RNAi worms was extracted from the whole worms in TRIzol (Life Technologies), following the manual instruction. The RNA samples were further analyzed by Qubit (Invitrogen) and Agilent Bioanalyzer. For each sample, ∼100ng RNA was used to generate the library using the Illumina TruSeq kit and sequenced in 150-bp paired reads using an Illumina MiSeq sequencer. Sequencing reads were mapped to the reference transcriptome, which contains sequencing data from TSA accession GDAG00000000.1 and additional NCBI deposited sequences. Reads were mapped with bowtie2 (Langmead & Salzberg, 2012) using default options other than allowing for multiple mapping (-k 100), due to the non-redundant nature of the transcript database. The mature and juvenile sexual *S. mediterranea* transcriptomes were generated as previously described (Xiang, Miller, Ross, Sanchez Alvarado, & Hawley, 2014). The CIW4 transcriptome was obtained from previous publication (Davies et al., 2017). Expression analysis was performed using DESEQ2 with default parameters (Love, Huber, & Anders, 2014).

For the genes expresses higher in sexually mature animals, comparing with the juvenile animals, the longest open reading frames were predicted from the RNAseq. Paircoil2 (McDonnell, Jiang, Keating, & Berger, 2006) with default parameters was used to predict the Coiled-coil domain and hmmscan (version 3.0rc2) (http://hmmer.org/) against PFAM-A (Finn et al., 2016) with a minimum e-value of 0.01 was used to predict the Zinc-finger domain.

Gene Ontology (GO) terms was analyzed for the genes decreased after *nhr-1* RNAi feeding 6 or 8 times (p<0.01). GO (Gene Ontology, 2015) were assigned to each *S. mediterranea* gene based on homologous PFAM domains and significant Swissprot hits. GO term enrichment was performed using the R package topGO (Alexa & Rahnenfuhrer, 2010).

### Molecular cloning and RNAi

Genes were cloned from the cDNA library, which was generated by reversed transcription from the mRNA of sexually matured *S. mediterranea*. The target genes were amplified with gene specific primers with overhangs and cloned to the pPR-T4P vectors (J. Rink) as previously described (Adler, Seidel, McKinney, & Sanchez Alvarado, 2014). DsRNA was produced in the bacteria and mixed with liver for RNAi feedings as previously described (Reddien, Bermange, Murfitt, Jennings, & Sanchez Alvarado, 2005).

### Whole mount DAPI and Lectin staining

Whole mount DAPI and PNA staining was done as previously described(Chong, Stary, Wang, & Newmark, 2011) with the following modifications. The sexual adult animal was killed in 10% N-acetyl cysteine (Sigma-Aldrich) in PBS for 5min. Then, the animal was fixed in 4% PFA, which containing 10% acetic acid for 1hr. After washout the fixation solution, the animal was treated in 10%SDS for 10min, which was followed by 1%NP40 0.5%SDS in PBS 20min. Bleach was done in 1.2%H2O2 with 5% formamide in 0.5xSSC above white light, which usually take 4hrs. After wash, the animal was blocked in gelatin Blocking buffer (0.6%BSA and 0.45%Fish gelatin in PBS) for 1hr and stained by staining solution (DAPI 5ug/ml and PNA-FITC 1:1000 in gelatin Blocking buffer) at 4°C for 2 days. After washing, the animal was fixed and mounted in 80% glycerol containing 2.5 % DABCO (Sigma-Aldrich) in PBS.

For screening, the sample was imaged by PerkinElmer Ultraview spinning-disc microscope and the max z projection was used to get the final image. For high resolution whole mount testes staining, which was used to identify the cell meiosis stages, the images were taken by LSM 510 META (Zeiss) with 100x oil lens and reconstructed by IMARIS.

### Telomere fluorescence *in situ* hybridization

Sexually mature planarians were killed in 7.5% N-acetyl cysteine (Sigma-Aldrich) in PBS for 5min, followed by a fixation in 4% Paraformaldehyde, Triton X-100 0.3% in PBS overnight at 4°C. After washing 3 times with PBS containing Tween-20 (0.3%), the sample was dehydrated through 30% sucrose and followed by embedding with OCT compound (Tissue-Tek, CA). Sagittal sections with 20µm thickness were cut using a Leica CM3050S cryostat (Leica Biosystems Inc. Buffalo Grove, IL).

Telomere probes (sequence: [TTAGGG]×7) were labeled with DIG-dUTP with Terminal Transferase (Roche) according to the manufacturer’s protocol. Labeled probes were suspended in deionized formamide (Sigma). For hybridization, a Master Hybridization Mix (4× Saline-Sodium citrate buffer [SSC], 20% dextran sulfate, 2 mg/mL nuclease-free Bovine Serum Albumin [BSA], 50% deionized formamide in ddH2O) was mixed with 1/10^th^ volume of telomere probes. DNA were denatured at 70 °C for 5 min. Hybridization was carried at RT for 36 to 48 hours. Slides were then washed with 2× SSC, 0.5× SSC, and TNT (100 mM Tris-HCl, 150 mM NaCl, 0.1% Tween-20) for 15 min each at RT. Anti-DIG-Rhodamine (Sigma Aldrich) was used at 1:200 in TNB buffer (5% Fetal Bovine Serum [FBS] in TNT) and incubated overnight. The following day the slides were washed with TNT buffer and stained with DAPI before imaging.

### Whole mount *in situ* hybridization

The RNA probe and in situ hybridization was done as previously described (Chong, Collins, Brubacher, Zarkower, & Newmark, 2013; King & Newmark, 2013; Wang, Zayas, Guo, & Newmark, 2007), with the following modifications. Animals were fixed in 4% formaldehyde in 1× PBS for 1hr. Animals were bleached in 3%H2O2 with 3% formamide in 0.5xSSC for 3-4hrs. The sample was mounted in Glycerol (80%), DABCO (2.5%) (Sigma-Aldrich) in PBS.

For fluorescent WISH, images were taken by Zeiss LSM-510 or PerkinElmer Ultraview spinning-disc microscope and Z projected, if not specified. For colorimetric WISH, images were taken by Leica M205 FA.

### Hematoxylin and eosin staining

The animals were killed in 5% N-acetyl cysteine (Sigma-Aldrich) in PBS for 5min, followed by a fixation in 4% Paraformaldehyde, Triton X-100 0.3% in PBS overnight at 4°C. After washing 3 times with PBS containing Tween-20 (0.3%), the sample was dehydrated, mounted in paraffin, serial sectioned (5µm thickness) and stained by H&E as previously describe (Adler et al., 2014).

The images were taken by Leica DM6000 with 20X objective lens and merged using Photoshop.

### Oil red O staining and quantification

For staining on sections, the sexual adult animal was killed in 7.5% N-acetyl cysteine (Sigma-Aldrich) in PBS for 5min and fixed in 4% paraformaldehyde, Triton X-100 0.1% in PBS overnight at 4°C and rinsed with 1X PBS for 3 times. Fixed worm was dehydrated through 30% sucrose and followed by embedding with OCT compound (Tissue-Tek, CA). Cryo sections with 14µm thickness were cut using a Leica CM3050S cryostat (Leica Biosystems Inc. Buffalo Grove, IL). After washing off OCT (3 × 5 min with PBS), cryo sections were pre-treated with propylene glycol for 2 min, followed by incubation with pre-warmed Oil Red O solution (EMS, cat# 26504-01) for 20min at 37°C. The slides were rinsed with 85% propylene glycol for 15 secs, and then with pre-warmed PBS at 37°C for 5 min twice. After DAPI (1:1000) stained at 4°C overnight, slide was mount and coversliped with ProLong Gold Antifade Mountant media (Thermo Fisher Scientific, cat# P36930). For whole mount samples, worm was pretreated by N-acetyl cysteine and fixed as above. After washing by 0.1% TirtonX-100 in PBS once and in PBS for additional two times, the worm was bleached in 3%H2O2 with 3% formamide in 0.5xSSC for 3-4hrs. After washing out the bleach solution, the worm was stained by Oil Red O with following modifications. Soaked in Oil Red O solution for 30min. After DAPI staining, the sample was post fixed with 4% Paraformaldehyde, Triton X-100 0.3% in PBS for 20min at RT. After wash, the worm was cleared in 80% glycerol at 4°C for 24 hrs and mounted in Glycerol (80%), DABCO (2.5%) (Sigma-Aldrich) in PBS.

Three channel transmitted light images of the planarian tail sections were acquired on an Olympus VS120 Slide Scanner and processed similar to previous work(O’Rourke, Soukas, Carr, & Ruvkun, 2009). Red, green and blue channels were separated, and an average of the green and blue was subtracted from the red. This image was thresholded to identify only regions of appreciable oil red staining and the red intensity in those regions was summed. The entire worm slice was segmented by thresholding to quantify its area. The image of whole mount sample was taken by Axioplan 206 std (Zeiss) with 10x lens.

### ACS and Acetyl-CoA treatment

Acyl-coenzyme A Synthetase (Sigma) was dissolved in nuclear free water to a final concentration of 0.25u/µl. After *nhr-1* RNAi feeding 4 times, ACS solution (2.5µl per worm per feeding) was mixed with *nhr-1* RNAi food (10µl per worm per feeding) for the following 6 feedings. Instead of the ACS solution, nuclease free water was mixed with *nhr-1* RNAi food at the same ratio for the last 6 feedings in the control animal. The present of testes and gland was tested 7 days after the last feeding.

Acetyl coenzyme A sodium salt (Sigma A2056) was used to rescue the *nhr-1* RNAi by feeding (30 nmol/worm, same protocol as above) or injection. After the sexually matured planarian had been fed by *nhr-1* RNAi food 4 times, injection was performed 3 days after each RNAi feeding for the following 6 feedings. Acetyl coenzyme A sodium salt was dissolved in nuclear free water to a final concentration of 25nmol/µl. 2µl Acetyl coenzyme A solution (or nuclease free water for the control group) was injected to every animal each time by using a Drummond Nanoject II microinjector (Broomall, PA)(Newmark, Reddien, Cebria, & Sanchez Alvarado, 2003). The present of testes and gland was tested 7 days after the last feeding.

### Statistical analyses

For statistical analyses of meiosis cells with bouquet in *nhr-1* RNAi animal vs. control animal, we used two-tailed Fisher’s exact test. Other statistical analyses were performed using Student’s t test.

## Accession Numbers

All RNA-seq datasets have been deposited in GEO: Accession number for *nhr-1*(*RNAi*) RNA-seq is GSE107756. Accession number for sexual fragment regeneration data is <u>GSE107869</u>.

## Supplementary figure legends

**Figure S1, related to Figure 1.**
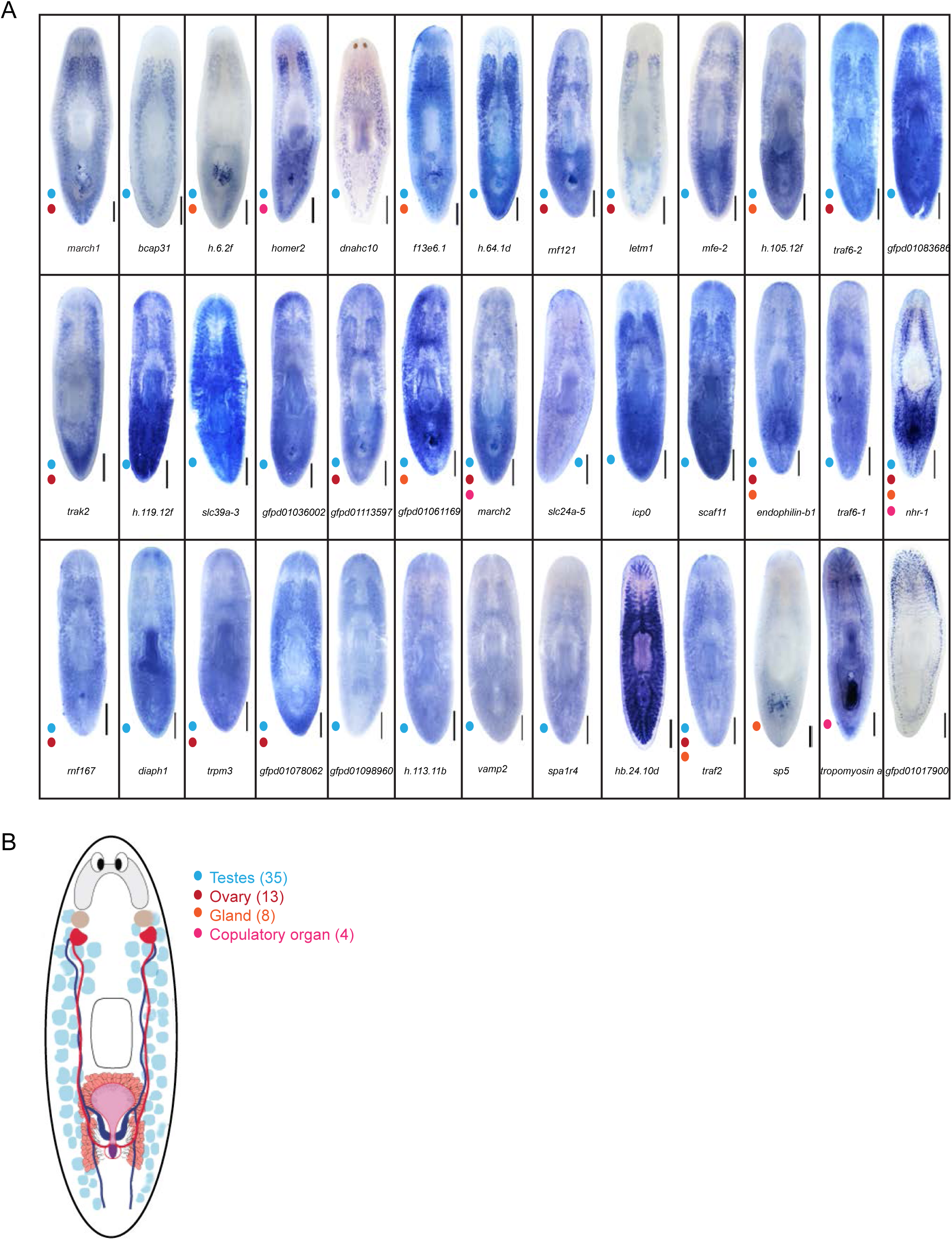
Majority of the adult worm enriched transcripts, which contain coiled-coil domain or zinc figure domain, express in the sexual reproductive system. F. WISH for candidate genes for screening. The color of dots represents different expression categories, which is explained in panel B. Scale bar: 750µm G. Summary of expression patterns of screened transcripts

**Figure S2, related to Figure 1.**
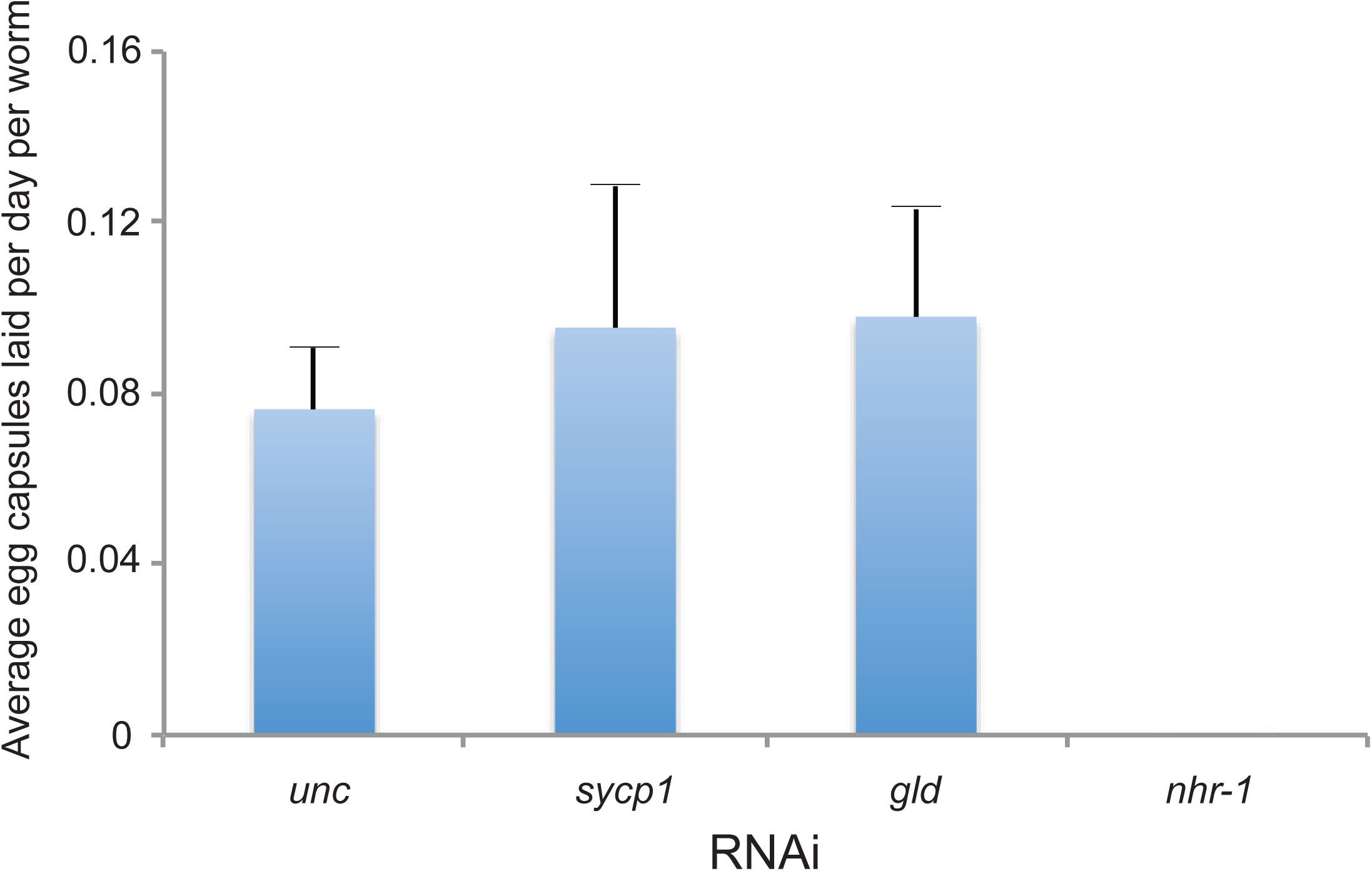
Histogram showing the cocoon number from different RNAi worms

**Figure S3, related to Figure 2.**
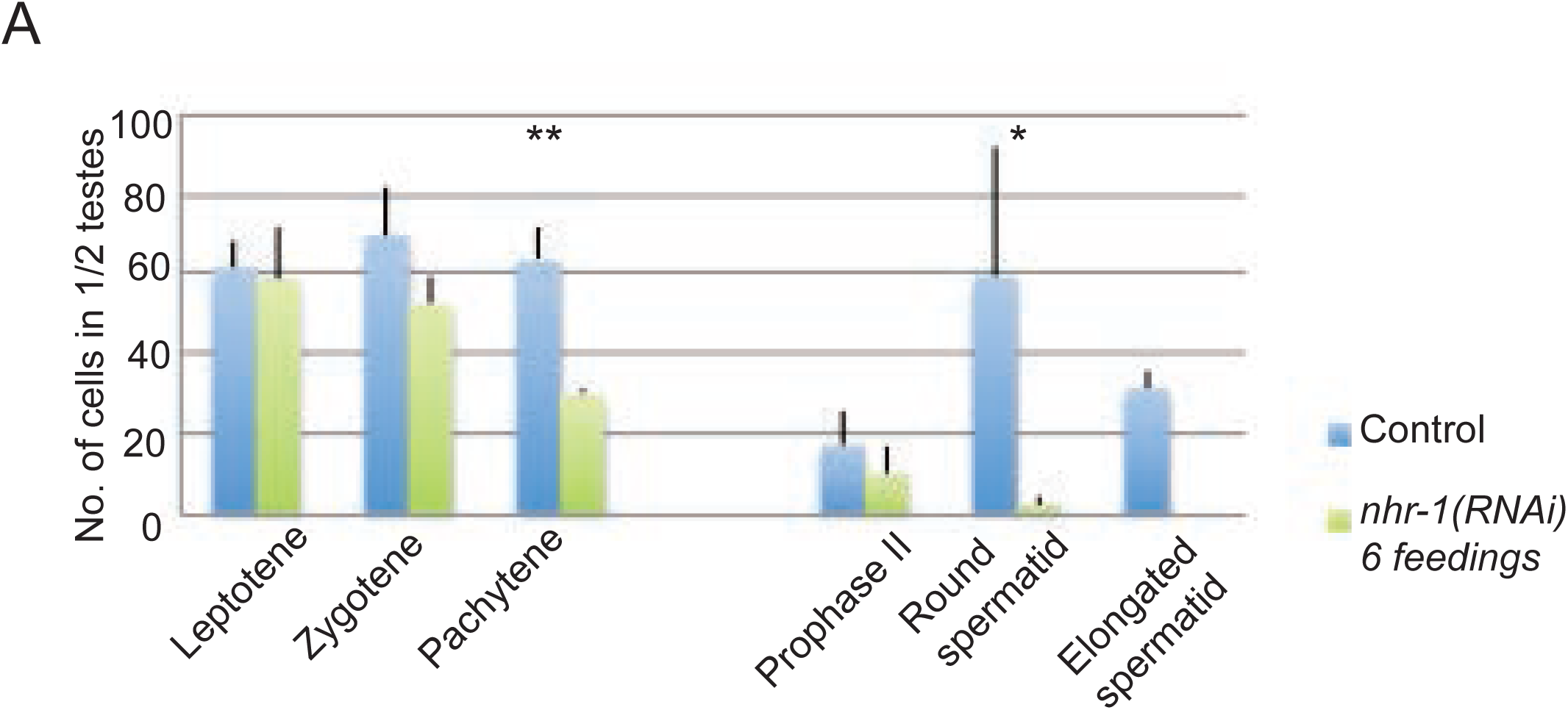
*nhr-1* RNAi decreased pachytene stage cells and telomere formation in testis. (A)Histogram of cell numbers in different meiosis stages in half testis (n=4, t-test,*p<0.05, **p<0.01). (B)Representative image showing normal bouquet in *nhr-1*(*RNAi*) animals. Scale bar: 5µm

**Figure S4, related to Figure 2.**
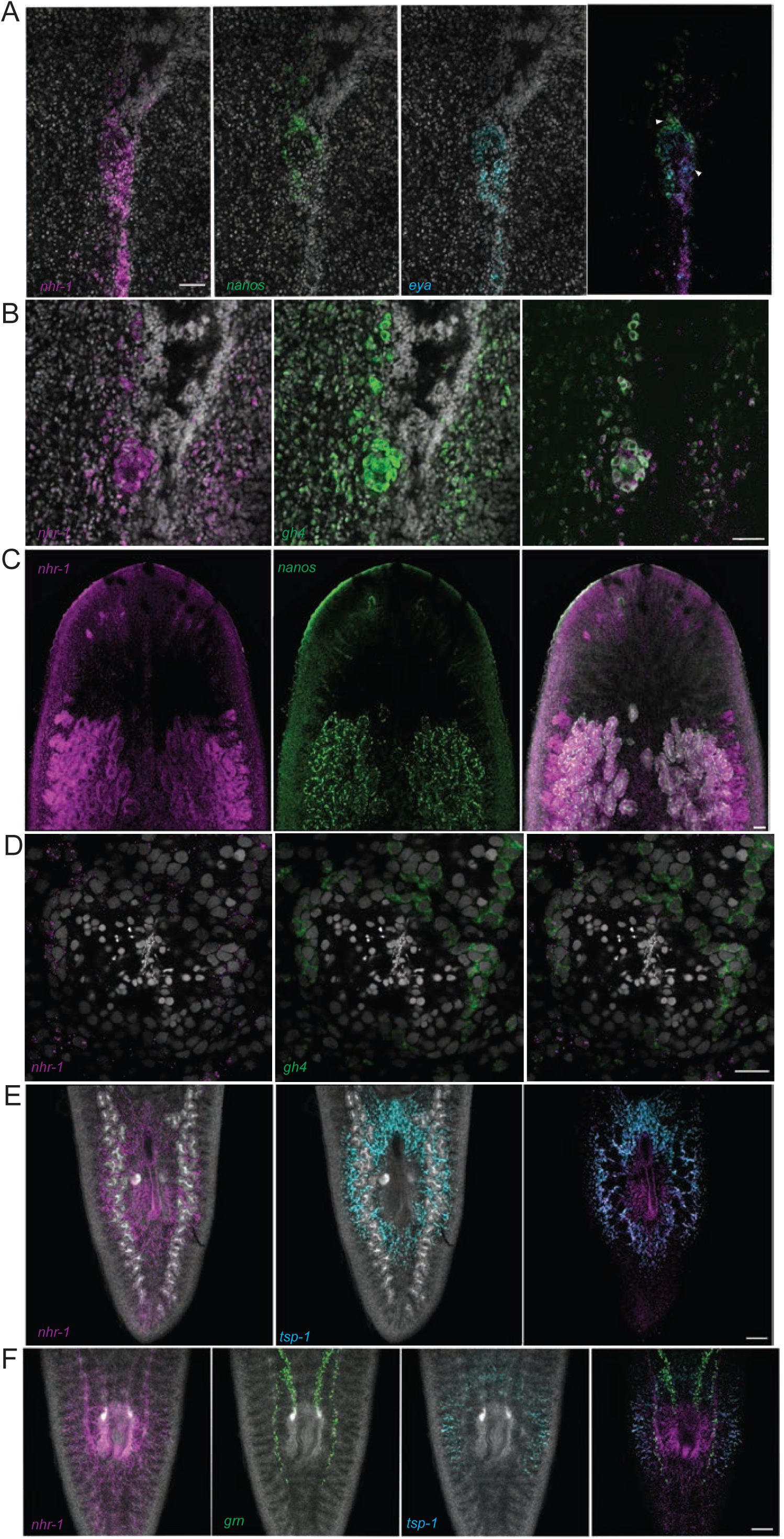
*nhr-1* expresses in most organs in the sexual reproductive system in adult planarian. (A)FISH for *nhr-1, nanos* and *eya* expression in ovary and oviduct. Scale bar: 20µm (B)FISH for *nhr-1* and *gh4* expression in ovary. Scale bar: 20µm (C)FISH for *nhr-1* and *nanos* in dorsal border region. *nanos* labels the germ stem cells in the testis. Scale bar: 100µm (D)FISH for *nhr-1* and *gh4* in testis. Scale bar: 20µm (E)FISH for *nhr-1* and *tsp-1* in dorsal tail. Scale bar: 100µm (F)FISH for *nhr-1, tsp-1* and *grn* in ventral tail. Scale bar: 100µm

**Figure S5, related to Figure 3.**
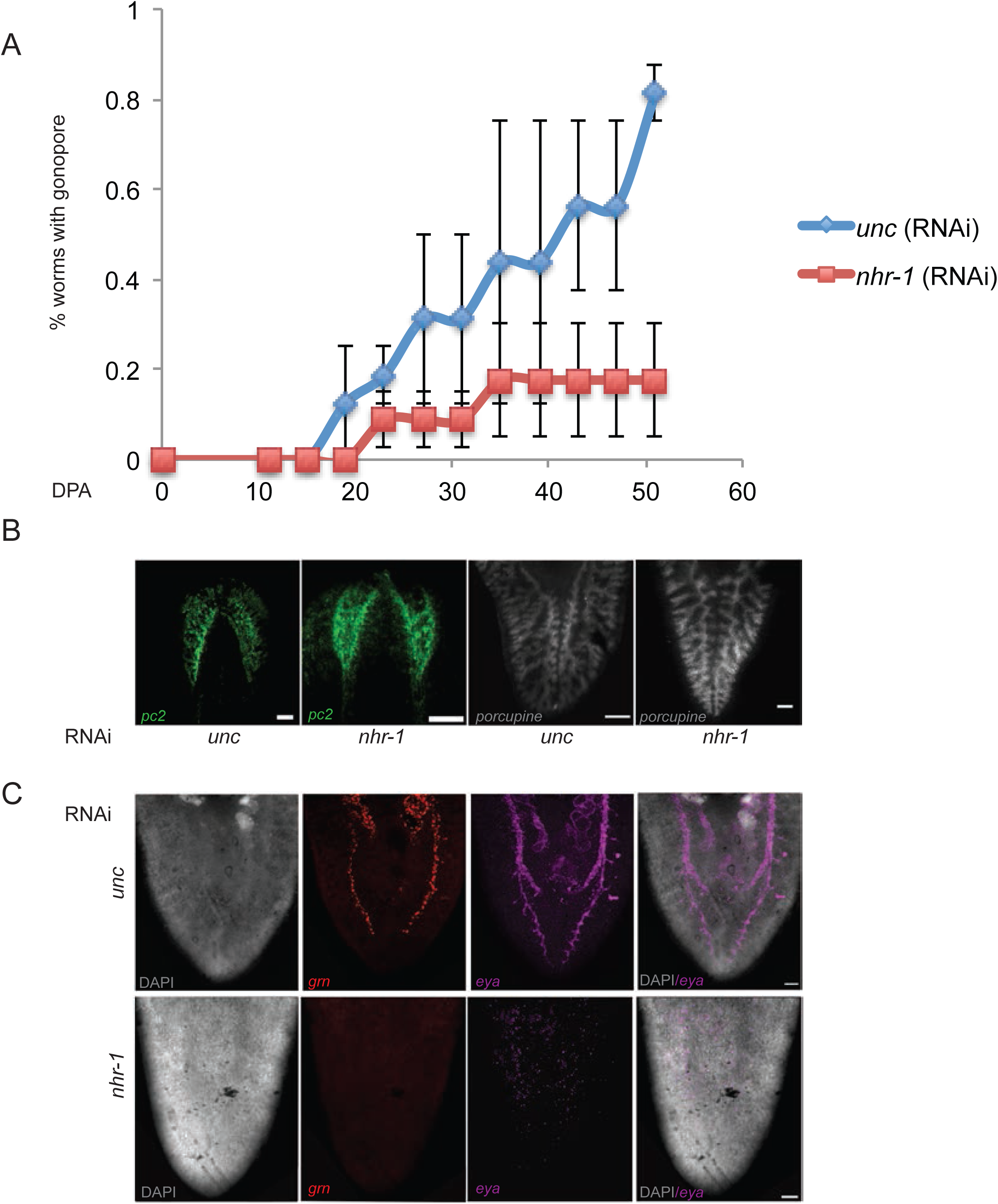
*nhr-1* is specifically required for reproductive system regeneration. (A)Percentage of worms with gonopore during regeneration. (B)FISH for *pc2* in regenerated head and *porcupine* in regenerated tail. Scale bar: 100µm (C)FISH for *grn* and *eya* in regenerated tail. Scale bar: 100µm

**Figure S6, related to Figure 5.**
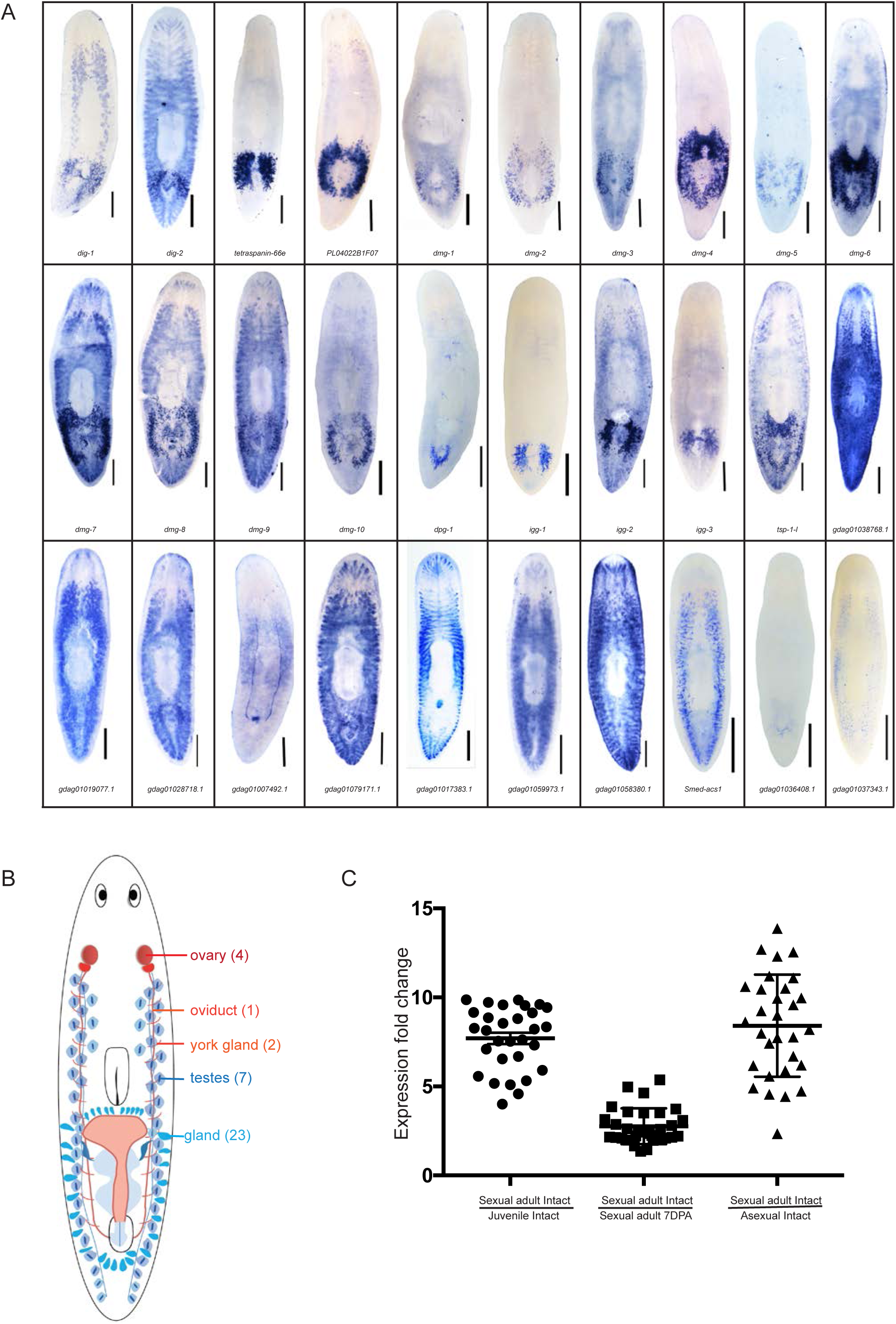
*nhr-1* downstream genes, which decreased after *nhr-1* RNAi feeding 4 times, are enriched in the reproductive system. (A)WISH of the 30 transcripts, which decreased after *nhr-1* RNAi feeding 4 times. Scale bar: 750µm (B)Summary of the 30 transcripts expression. (C)Expression fold changes of genes affected after 4 rounds of *nhr-1* RNAi feeding in asexual, juvenile, decapitated sexual adult fragments (D0) and 7 days post amputation (D7). Each dot shows ratio of single transcripts. Mean ±SEM of each data group showed by lines in the figure. n=3 for juvenile, sexual adult and sexual adult fragments, each containing 3 worms. n=4 for asexual worms as previously reported (Davies et al., 2017).

**Figure S7, related to Figure 5.**
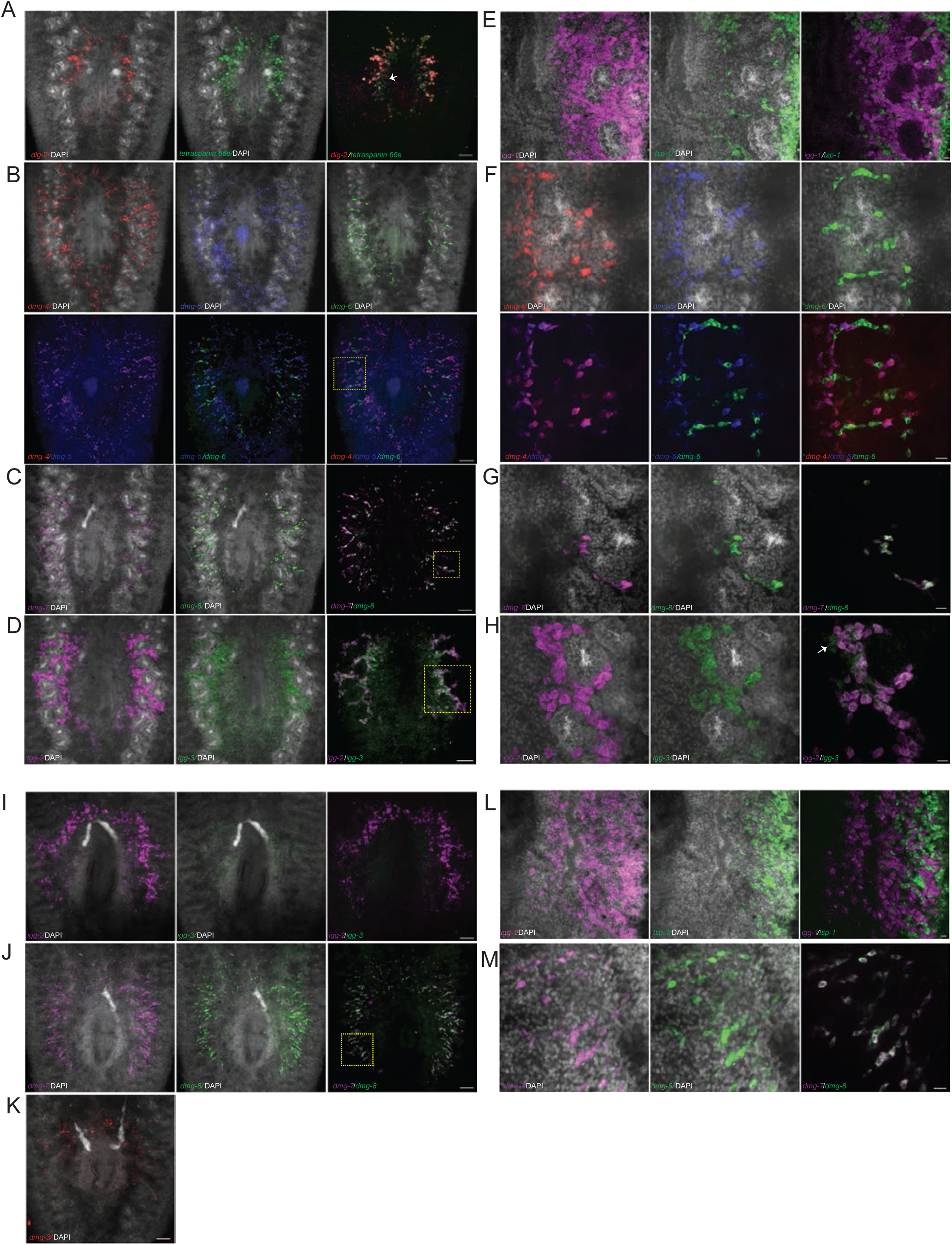
*nhr-1* downstream genes, which decreased after *nhr-1 RNAi* feeding 4 times, express in different regions of the glands in sexual mature planarian. Panel A to H are the in the dorsal tail region. Panel I to M are in the ventral tail region. (A)FISH for dorsal expression of *dig-2* and *tetraspanin 66e*, which express in region 1. Scale bar: 100µm (B)FISH for dorsal expression of *dmg-4, dmg-5* and *dmg-6*, which express in region 2. The yellow box shows the region for further zoom in in panel F. Scale bar: 100µm (C)FISH for dorsal expression of *dmg-7* and *dmg-8*, which express in region 2. The yellow box shows the region for further zoom in in panel G. Scale bar: 100µm (D)FISH for dorsal expression of *igg-2* and *igg-3*, which express in region 4. The yellow box shows the region for further zoom in in panel H. Scale bar: 100µm (E)FISH for dorsal expression of *igg-1* and *tsp-1* in single cell layer. Scale bar: 20µm (F)FISH for dorsal expression of *dmg-4, dmg-5* and *dmg-6* in single cell layer. Scale bar: 20µm (G)FISH for dorsal expression of *dmg-7* and *dmg-8* in single cell layer. Scale bar: 20µm (H)FISH for dorsal expression of *igg-2* and *igg-3* in single cell layer. Scale bar: 20µm (I)FISH for ventral expression of *igg-2,* which expresses in region 2, and *ig3*. Scale bar: 100µm (J)FISH for ventral expression of *dmg-7* and *dmg-8*, which express in region 3. The yellow box shows the region for further zoom in in panel M. Scale bar: 100µm (K)FISH for ventral expression of *dmg-3*, which expresses in region 1. Scale bar: 100µm (L)FISH for ventral expression of *igg-1* and *tsp-1* in single cell layer. Scale bar: 20µm (M)FISH for dorsal expression of *dmg-7* and *dmg-8* in single cell layer. Scale bar: 20µm

**Figure S8, related to Figure 5.**
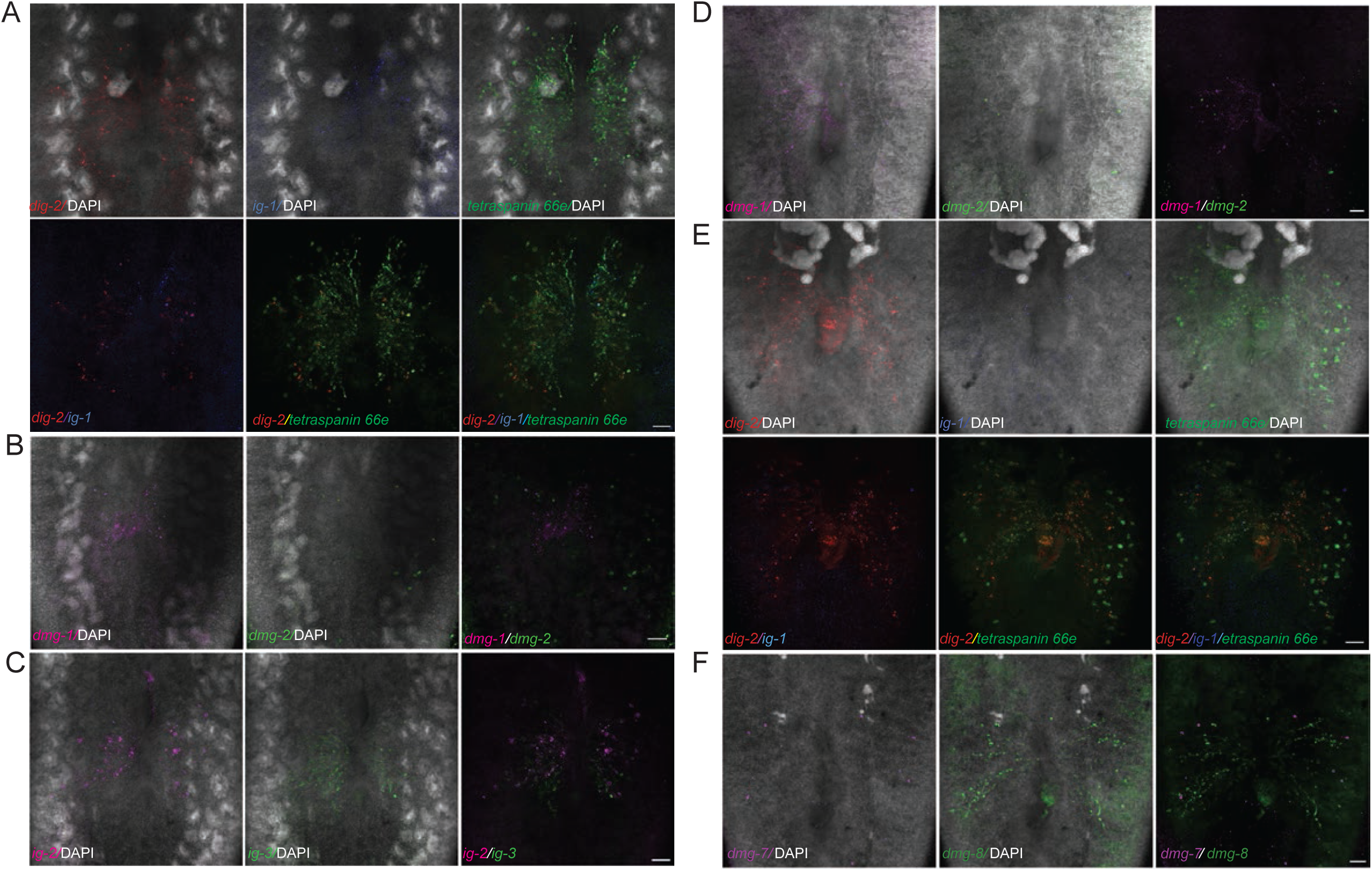
*nhr-1* is essential for the expression of novel gland genes, which decreased after *nhr-1 RNAi* feeding 4 times. This figure shows the representative images of FISH for expression of novel gland genes after *nhr-1* RNAi feeding 4 times in adult planarian. Panel A to C show the image in the dorsal region, while panel D to F show the image in the ventral region. (A)FISH for expression of *dig-2, igg-1* and *tetraspanin 66e* in dorsal tail region. Scale bar: 100µm (B)FISH for expression of *dmg-1* and *dmg-2* in dorsal tail region. Scale bar: 100µm (C)FISH for expression of *igg-2* and *igg-3* in dorsal tail region. Scale bar: 100µm (D)FISH for expression of *dmg-1* and *dmg-2* in ventral tail region. Scale bar: 100µm (E)FISH for expression of *dig-2, igg-1* and *tetraspanin 66e* in ventral tail region. Scale bar: 100µm (F)FISH for expression of *dmg-7* and *dmg-8* in ventral tail region. Scale bar: 100µm

**Figure S9, related to Figure 6.**
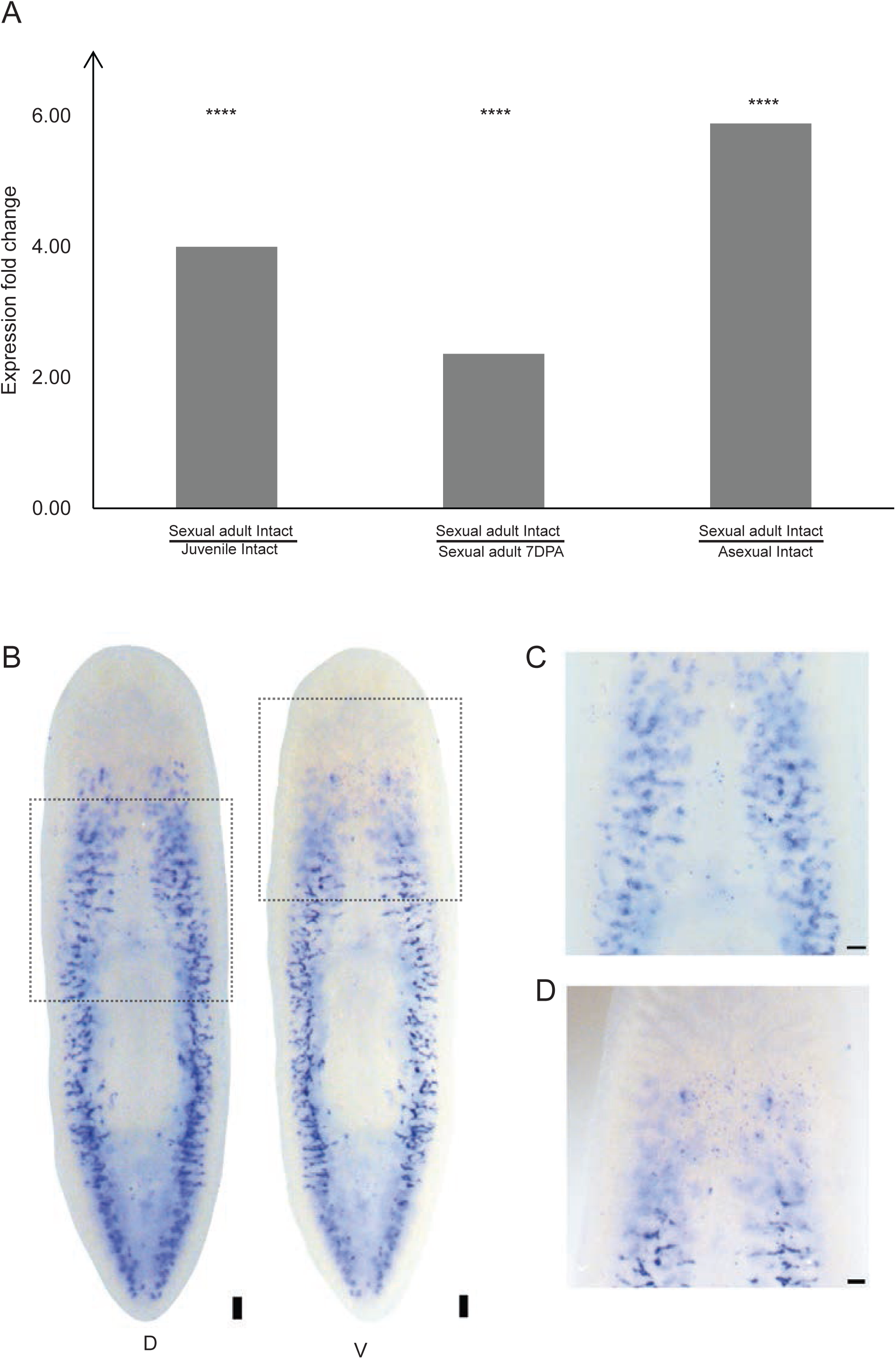
*Smed-acs-1* expression in sexual adult animal. (A)*Smed-acs-1* expression in intact asexual, juvenile, decapitated sexual adult (D0) and sexual adult fragments (7 days after amputation, D7) according to RNAseq data. Dot shows RPKM value in each replicate. Bar graph indicates the Mean±SEM value for each sample. n=3 for juvenile, sexual adult and sexual adult fragments, each containing 3 worms. n=4 for asexual worms, as previously reported (Davies et al., 2017). *represents the adjusted p value for each comparison group. ****p<0.0001. (B)WISH of *Smed-acs-1* in dorsal and ventral region. The dorsal region in the white square was further zoom in in the panel B. The ventral region in the white square was further zoom in in the panel C. Scale bar: 100µm (C)Zoom in of *Smed-acs-1* dorsal expression posterior of neck region. Scale bar: 50µm (D)Zoom in of *Smed-acs-1* ventral expression in the ovary region. Scale bar: 50µm

**Figure S10, related to Figure 7.**
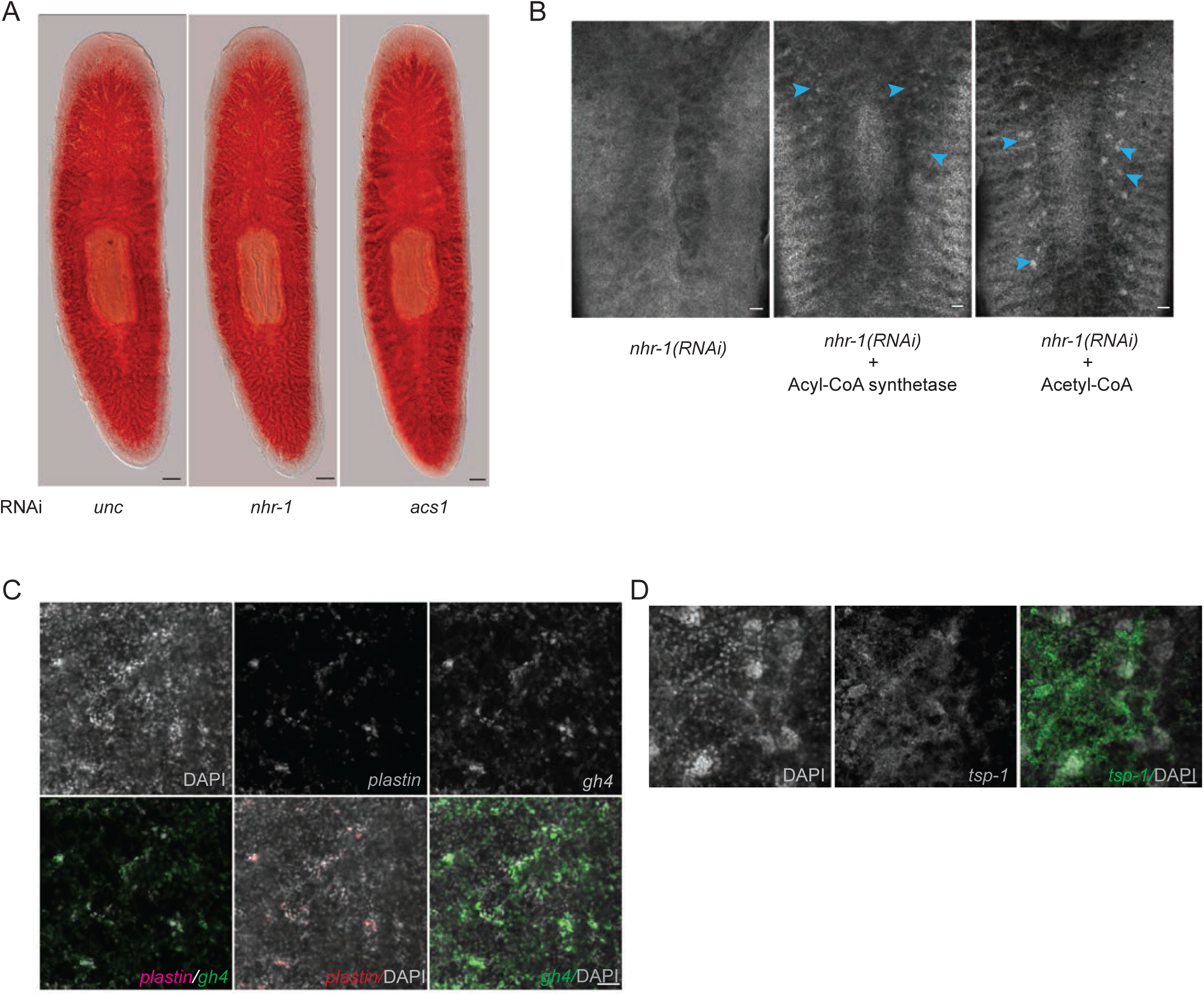
*nhr-1* maintains reproductive system through lipid metabolic pathway in sexual planarian. (A)Oil Red O staining of asexual animal. Scale bar: 500µm (B)DAPI staining in the dorsal tail region of the sexual mature animal after long term *nhr-1* RNAi. Blue arrowheads point to the testes after rescue, which enrich DAPI signal. Scale bar: 100µm (C)FISH of *plastin* and *gh4* in dorsal testes region after Acetyl-CoA rescue. Scale bar: 50µm. (D)FISH of *tsp-1* in dorsal tail region after Acyl-CoA rescue. Scale bar: 20µm.

**Movie-S1**

3D reconstruction of nuclei in half testis with whole-mount DAPI staining in normal sexual adult planarian.

**Movie-S2**

3D reconstruction of nuclei in half testis with whole-mount DAPI staining in sexual adult planarian after *nhr-1* RNAi feeding 6 times

**Table S1: related to Figure 1**

Expression level changes of the 34 genes identified to be expressed higher in decapitated adult fragments (D0) when compared to juvenile animals. D7 shows decapitated sexual adult fragments 7 days post amputation.

**Table S2: related to Figure 4**

Expression level changes of the 30 genes down-regulated after 4 rounds of *nhr-1*(RNAi) feeding. D0 and D7 show decapitated sexual adult fragments at 0 and 7 days post amputation.

**Table S3: related to Figure 1 and Figure 5**

Primers used for gene cloning

